# Bioactive Enhanced Adjuvant Chemokine Oligonucleotide Nanoparticles (BEACON) for Mucosal Vaccination Against Genital Herpes

**DOI:** 10.1101/2025.07.31.667899

**Authors:** Sachin H. Bhagchandani, Stephen Ehrenzeller, Ivan S. Pires, Namit Chaudhary, Carmen J. Booth, Dong-il Kwon, Charalampia Koutsioumpa, Christopher A. Baker, Claire Laxton, Keyla Santos Guedes de Sá, Cullen Matthews, Priya Gill, Sarah Li, Anna Olszowka, Andrew Hudak, Suzanne Fischer, Rafael Bayarri-Olmos, William Brenham Hooper, Akiko Iwasaki

## Abstract

Genital herpes, caused by herpes simplex virus-2 (HSV-2), remains a prevalent sexually transmitted infection with no available vaccine. Effective local immunity—including tissue-resident memory T cells (T_RMs_) and luminal antibodies—provides immediate viral control. Here, we developed Bioactive Enhanced Adjuvant Chemokine Oligonucleotide Nanoparticles (BEACON) formed via electrostatic interactions between CpG oligodeoxynucleotides (CpG ODN) and the chemokine CXCL9. This adjuvant enhances antigen-presenting cell engagement and innate immune signaling, promotes CD8^+^ T cell recruitment, and reduces local neutrophilic inflammation relative to CpG ODN. Co-administered vaginally with HSV-2 glycoproteins following intramuscular priming, BEACON improved protection against HSV-2 by increasing local CD8^+^ T_RM_ populations and mucosal IgG and IgA responses. Vaccine-mediated protection required local delivery of both antigen and adjuvant, and was significantly reduced by CD8^+^ T cell or B cell depletion. These findings highlight the potential of engineered mucosal adjuvants for vaccines targeting genital herpes and other sexually transmitted infections.

**One-sentence summary:** We developed BEACON, a mucosal adjuvant that elicits protective immunity against genital herpes simplex virus 2 infection in mice.

## INTRODUCTION

Genital herpes, a widespread sexually-transmitted infection caused primarily by herpes simplex virus type 2 (HSV-2), affects over half a billion individuals worldwide (*1*). HSV-2 establishes lifelong latency in dorsal root ganglia (DRG), with periodic reactivation, recurrent symptoms, and viral shedding, even in asymptomatic individuals (*2*, *3*). HSV-2 infection also increases susceptibility to human immunodeficiency virus 1 (HIV-1), compounding the global health burden (*4–6*). There are no approved vaccines, and current antiviral therapies mitigate symptoms without preventing transmission or reactivation (*7*). While parenteral vaccination with recombinant HSV-2 glycoproteins (gB and/or gD) failed to confer robust protection (*8–10*), efforts to develop more effective vaccines have been hindered by both viral immune evasion mechanisms and the lack of robust mucosal immunity at the site of entry. Messenger RNA-based platforms utilizing lipid nanoparticles (LNPs) have been tested, but still face significant challenges in eliciting durable local immunity at mucosal surfaces (*11*, *12*).

A major limitation of conventional parenteral HSV-2 vaccine approaches is their failure to establish a robust population of CD8^+^ tissue-resident memory T (T_RM_) cells and local antibody responses at the vaginal mucosa, the site of viral entry (*13–15*). We previously introduced a “prime and pull” vaccination strategy, in which topical chemokines were introduced after systemic priming to promote local recruitment of effector CD8^+^ T cells (*16*). Although this approach offers partial protection, it does not engage B cell-mediated immunity, which is essential for neutralizing the virus at the mucosal surface (*17–20*). Furthermore, local CD8^+^ T_RM_ cells alone cannot prevent primary HSV infection or reduce viral titers (*16*). Local administration of high-dose intravaginal CpG oligodeoxynucleotides (CpG ODN), synthetic Toll-like receptor 9 (TLR9) agonists, reduced viral loads in preclinical models, but elicited substantial vaginal inflammation (*21–24*). A key challenge is to develop a vaccine strategy that simultaneously recruits effective tissue-resident T cells and elicits robust B cell responses without causing excessive local inflammation.

While several intravaginal platforms, including replication-defective HSV-2 vaccines and viral vector-based approaches, have shown promise in murine models, these strategies have not translated into clinical success (*25–29*). One limitation is the inherently tolerogenic nature of the vaginal mucosa, which can suppress immune priming, diminish the efficacy of local immunization and promote local inflammation (*25*, *30–33*). Alternatively, systemically primed immunity can be leveraged to generate appropriate mucosal responses, for example, intramuscular priming to establish a pool of antigen-specific T and B-cells, followed by a mucosal boost to drive their recruitment to and activation at the site of infection (*34*). Achieving this requires a booster that safely enhances immune activation in the vaginal mucosa without causing tissue damage or inflammation (*35*).

To address these challenges, we developed a mucosal adjuvant that combines CpG ODN and CXCL9 to generate Bioactive Enhanced Adjuvant Chemokine Oligonucleotide Nanoparticles (BEACON). Here, we examine the use of BEACON as both an innate antiviral agent and a vaccine adjuvant against genital HSV-2 infection in mice, and probe its protective mechanisms.

## RESULTS

### Development of Bioactive Enhanced Adjuvant Chemokine Oligonucleotide Nanoparticles (BEACON)

Although vaginal application of CXCL9 or CXCL10 following systemic priming facilitates CD8^+^ T cell recruitment to the genital mucosa and enhances protection against wild-type (WT) HSV-2 (*16*, *36*), this approach does not significantly reduce viral titers (*16*, *18*, *20*). Similarly, high doses of intravaginally-administered CpG ODN mediate potent antiviral effects, albeit with substantial vaginal inflammation (**fig. S1A-B**) (*21*). Reducing the CpG ODN dose alleviates inflammation, but also compromises its protective efficacy (**fig. S1C-D**).

To improve the delivery and efficacy of CpG ODN, we formulated BEACON based on electrostatic interactions between negatively charged CpG ODN and the cationic chemokine CXCL9 (**Fig. 1A**). This approach yielded stable, uniformly sized particles (150-200 nm), as assessed by dynamic light scattering (**Fig. 1B; fig. S2A**), and confirmed morphologically by negative-stain and cryogenic transmission electron microscopy (**Fig. 1C; fig. S2B**). Equivalent formulations were generated using human CXCL9 and CpG 1018, the latter a clinically approved TLR9 agonist used in hepatitis B vaccines (**fig. S2C**). Optimization of nanoparticle assembly required careful modulation of the CpG-ODN-to-chemokine molar ratio (**fig. S2D**). Stable complexes were formed in the presence of excess CXCL9, suggesting that CpG ODN can effectively crosslink multiple CXCL9 molecules (**fig. S2D**). Given the similar net charge per molecule, the resulting CXCL9-CpG ODN assemblies exhibited a slightly negative zeta potential, a physicochemical property favorable for in vivo administration (**fig. S2D**)(*37*). To maximize CpG ODN incorporation while avoiding the need for post-assembly purification, we minimized CpG input and identified an optimal molar ratio to obtain stable particles approaching the saturation threshold, as indicated by a plateau in zeta potential measurements (**fig. S2D**)(*38*).

**Figure 1.**
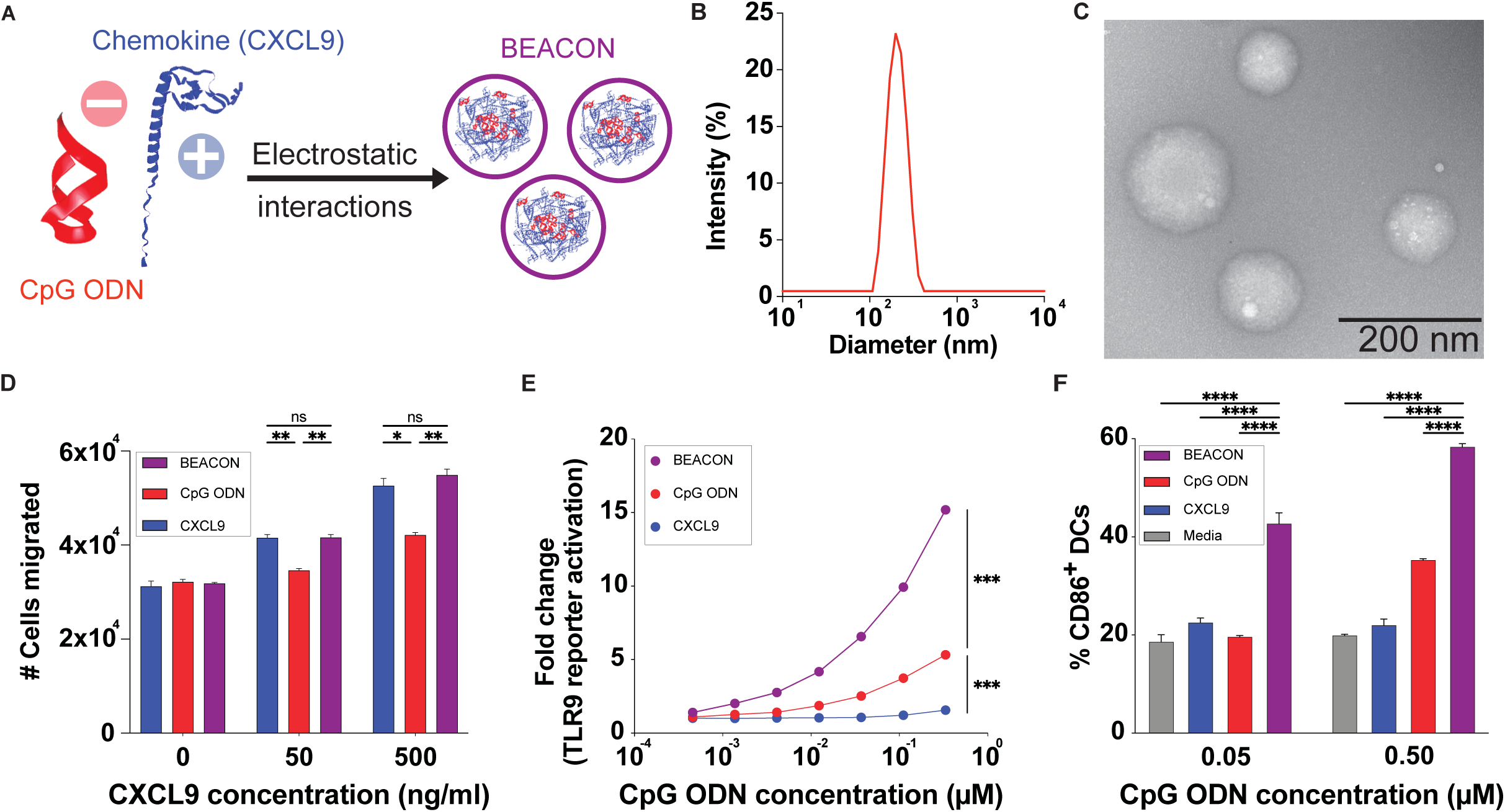
Design and characterization of Bioactive Enhanced Adjuvant Chemokine CXCL9 and CpG Oligonucleotide (CpG ODN) Nanoparticles (BEACON). (**A**) Schematic of BEACON resulting from electrostatic interactions between CpG ODN and CXCL9. (**B**) Hydrodynamic diameter of a mouse BEACON measured by dynamic light scattering. (**C**) Representative negative stain transmission electron microscopy images showing spherical morphology of mouse BEACON; scale bar, 200 nm. (**D**) Measurement of CXCL9 functional activity via chemotaxis assay, quantifying the number of T cells migrated across the Transwell^®^ in response to soluble CXCL9, free CpG ODN, or BEACON. The number of cells migrated was determined by Calcein AM fluorescence. (**E**) TLR9 signaling assessed by measuring fold change (relative to saline) in a HEK-mouse-TLR9 reporter assay 48 hours after treatment with free CpG ODN, CXCL9, or BEACON at the CpG concentrations indicated; CXCL9 alone was included as a negative control. (**F**) Frequency of CD86^+^ BMDCs detected after stimulation with BEACON compared to controls (free CpG-ODN, CXCL9, or medium alone). Data are plotted as mean ± SEM. Statistical significance in panels D - F was determined by two-way ANOVA with Tukey’s multiple-comparisons test: *P < 0.05, **P < 0.01, ***P < 0.001, ****P < 0.0001; ns, not significant.

Attempts to generate BEACON with CXCL10 yielded unstable aggregates (**fig. S2E-G**). We hypothesized that CXCL9’s extended, cationic C-terminal domain (amino acids 74-103) facilitates stable nanoparticle assembly (*39*). We found that deletion of this region abrogated particle formation (**fig. S2H**), while appending it to CXCL10 facilitated nanoparticle generation with consistent morphology (**fig. S2I**). These data highlight the critical role of the CXCL9 C-terminal motif in mediating stable CpG ODN packaging and underscore the potential of chemokine engineering for immunostimulatory nanoparticle design.

BEACON also retained chemoattractant function (**Fig. 1D**) and led to a three-fold increase in TLR9-mediated reporter activity compared to free CpG ODN alone (**Fig. 1E**), suggesting enhanced endosomal delivery. To assess functional activation of APCs, we incubated murine bone marrow-derived dendritic cells (BMDCs) with BEACON, CpG ODN, CXCL9, or medium alone and measured surface expression of the DC activation marker CD86. BEACON-treated BMDCs exhibited significantly higher frequencies of CD86⁺ DCs than cells treated with CpG ODN, CXCL9, or medium alone (**Fig. 1F**). These findings suggest the potential of BEACON as a dual-function platform capable of TLR9-mediated innate immune activation and CXCL9-directed lymphocyte recruitment with favorable biophysical and functional properties for mucosal immunization.

### BEACON targets CpG to APCs and limits local pathology

To determine whether BEACON could limit CpG-mediated mucosal inflammation, we administered low-dose CpG ODN (10 µg) alone or as BEACON intravaginally into Depo-Provera-treated WT mice (**Fig. 2A**). Mice treated with high-dose CpG ODN (100 µg) served as a positive control for vaginal inflammation. Histopathological analysis of vaginal tissue 48 hours post-administration revealed pronounced neutrophilic infiltration and epithelial thickening in the high-dose CpG ODN group (**Fig. 2B**). While inflammation was partially reduced by reducing the dose of CpG ODN, mice receiving the same dose delivered as BEACON exhibited markedly fewer histopathological features of inflammation (**Fig. 2C**) and fewer neutrophils in the vaginal tract (**fig. S3A-B**). We hypothesized that formulating CpG ODN into nanoscale BEACON could bias CpG delivery toward APCs, increase the valency of CpG within endosomal compartments, and limit recruitment of inflammatory neutrophils (*40*, *41*).

**Figure 2.**
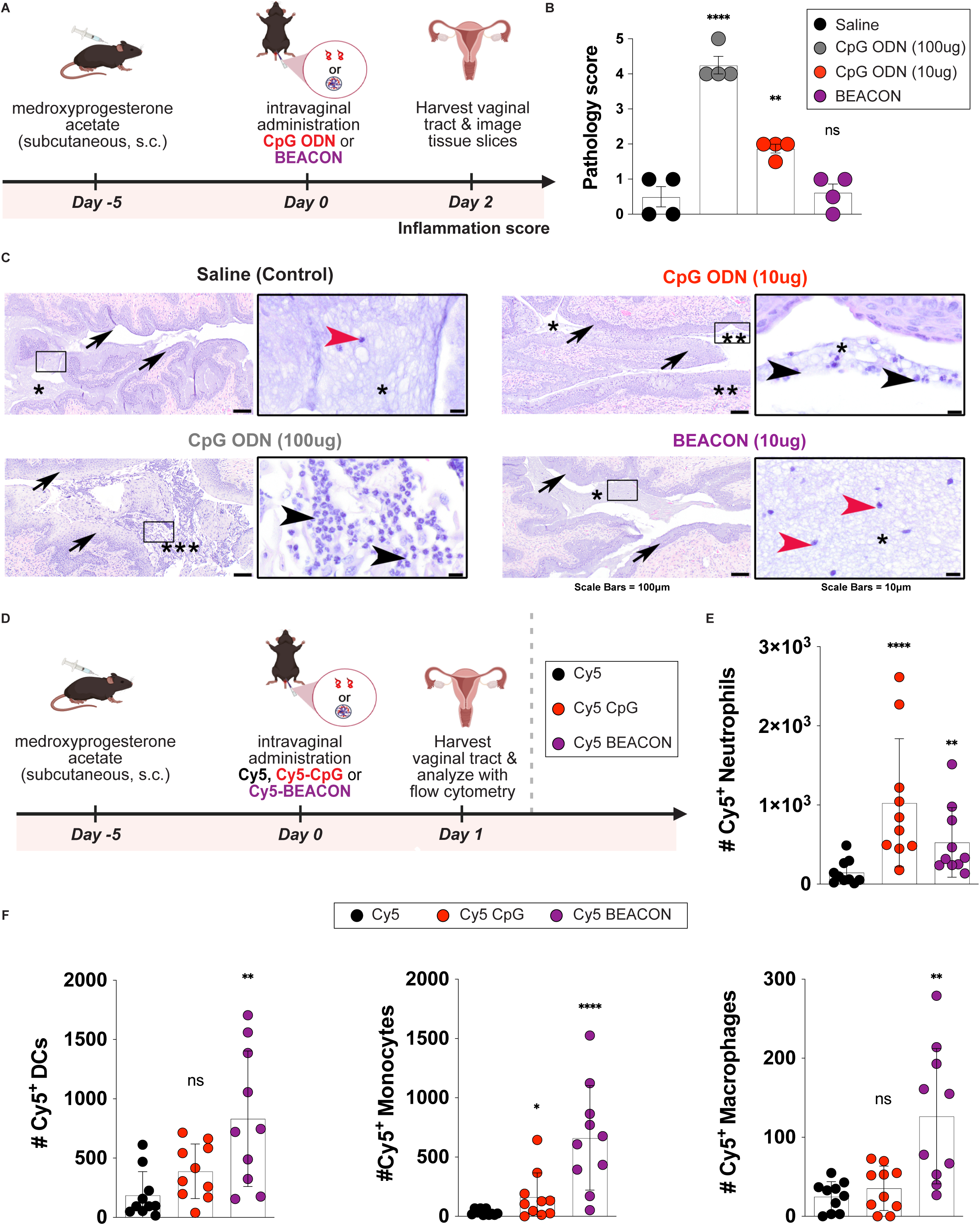
Intravaginally administered BEACON reduces CpG-associated mucosal inflammation and improves CpG delivery to vaginal APCs. **(A)** Experimental schematic for vaginal toxicity assessment. Depo-Provera pre-treated female C57BL/6 mice received intravaginal saline, CpG ODN (10 µg), BEACON (10 µg), or high-dose free CpG ODN (100 µg) as a positive control for inflammation. **(B)** Blinded histopathology score of vaginal tissues harvested 48 h after intravaginal administration. Each dot represents one mouse, n=4 per group. **(C)** Representative hematoxylin and eosin (HE)-stained vaginal sections collected 48 h after administration of CpG ODN, BEACON, or saline control. Insets indicate the field featured in the higher magnification images to the right of each panel. Scale bars are as indicated. **(D)** Experimental schematic for cellular uptake study using Cy5 control, Cy5-CpG, or Cy5-BEACON delivered intravaginally to Depo-Provera pre-treated female C57BL/6 mice. **(E)** Quantification of Cy5⁺ neutrophils in vaginal tissue by flow cytometry. **(F)** Quantification of Cy5⁺ APC subsets (DCs, monocytes, macrophages) in vaginal tissue. Each dot represents one mouse; data were pooled from two independent experiments (n=10 per group overall). Bars indicate mean ± SEM. Statistical significance in panels B, E, and F was determined by Kruskal-Wallis with Conover-Iman multiple comparisons procedure using Benjamini-Hochberg for controlling false discovery rate. *P < 0.05, **P < 0.01, ***P < 0.001, ****P < 0.0001; ns, not significant.

To assess whether reduced inflammation was associated with altered cellular targeting, we compared the biodistribution of fluorescently-labeled Cy5-BEACON to Cy5-labeled free CpG ODN (**Fig. 2D, fig. S3C**). BEACON demonstrated a ∼10-fold increase in uptake by APCs and a ∼two-fold reduction in neutrophil uptake (**Fig. 2E-F**). Our findings highlight the dual benefits of BEACON: reducing vaginal inflammation and optimizing CpG ODN delivery to APCs, enhancing immunomodulation function.

### BEACON recruits and retains CD8^+^ T cells and confers protection against genital HSV-2

We next evaluated whether BEACON can enhance mucosal recruitment and retention of HSV-specific CD8^+^ T cells using the established “prime and pull” approach (*16*). Mice received an adoptive transfer of gB-specific (gBT-I) CD8^+^ T cells before subcutaneous immunization with attenuated thymidine kinase (TK)-deficient HSV-2 (*42*), followed by intravaginal administration of saline, CpG ODN, CXCL9, or BEACON (**Fig. 3A, fig. S4A**). One day after treatment, BEACON-treated mice had nearly ten times more gBT-I CD8^+^ T cells in the vaginal mucosa than saline controls and twice as many as those treated with CpG ODN or CXCL9 alone (**Fig. 3B-C, fig. S4B-E**), with persistent elevation at four weeks post-administration (**Fig. 3D, fig. S4A**).

**Figure 3.**
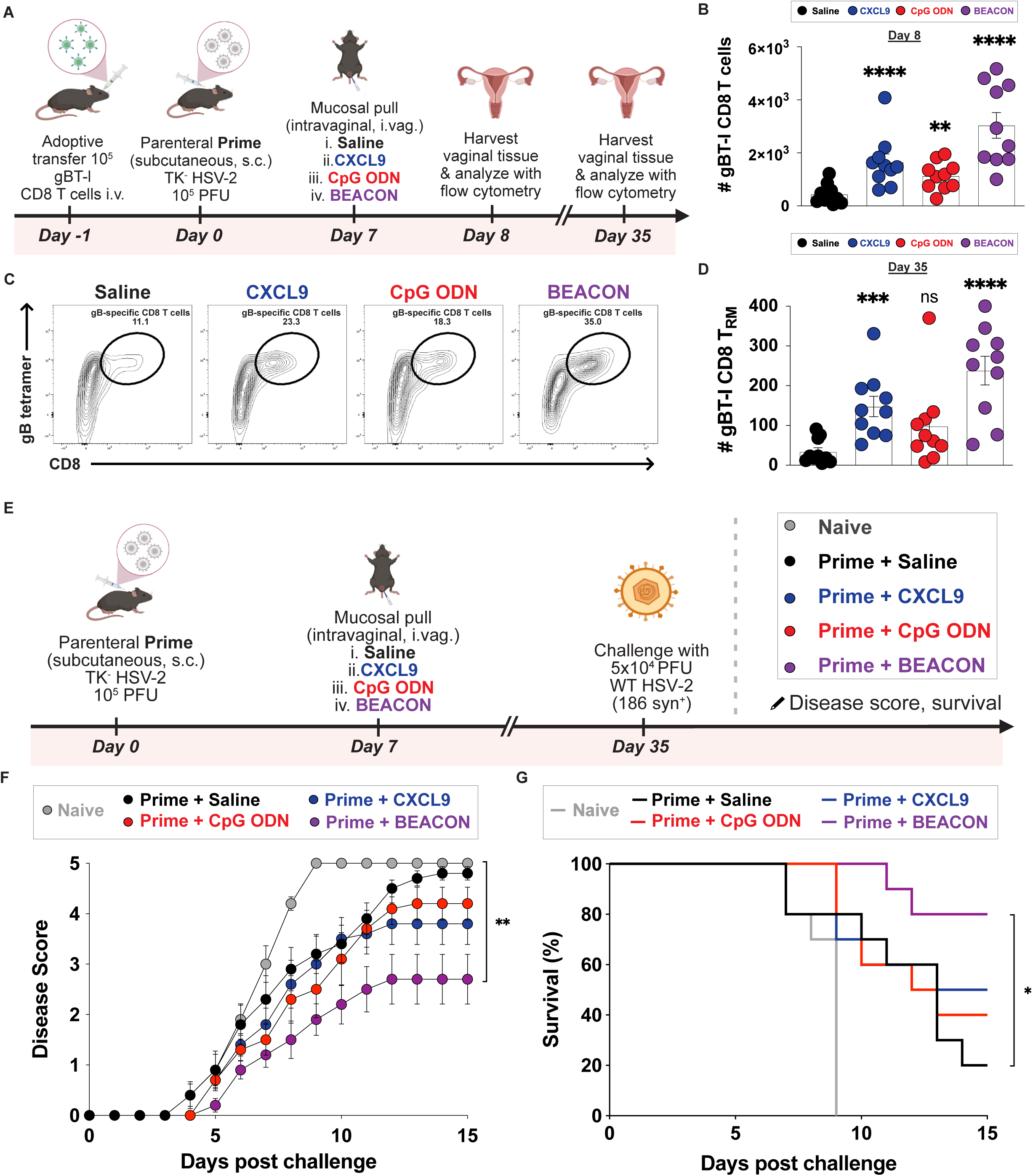
BEACON enhances recruitment and retention of HSV-specific CD8^+^ T cells in the vaginal tract and improves protection against WT HSV-2 challenge. **(A)** Experimental schematic. CD45.1^+^ gBT-I CD8^+^ T cells were adoptively transferred into recipient C57BL/6 mice, followed by subcutaneous priming with TK-deficient HSV-2 and intravaginal “pulling” with saline, CXCL9, CpG ODN, or BEACON at indicated time points. Vaginal tissues were harvested and analyzed on day 8 (acute recruitment) and day 35 (memory). **(B)** Number of gBT-I CD8^+^ T cells in the vaginal tract (day 8). **(C)** Representative flow cytometry plots for gB tetramer^+^ CD8^+^ T cells, gated on total T cells. **(D)** Quantification of gBT-I CD8^+^ T_RM_ (CD69^+^ CD103^+^) cells in the vaginal tract (day 35). **(E)** Experimental schematic for the challenge study using the same prime and pull regimen followed by WT HSV-2 challenge on day 35. **(F)** Clinical disease scores and **(G)** Kaplan-Meier survival curves following intravaginal challenge with WT HSV-2. Data in bar graphs are presented as mean ± SEM, with individual points representing one mouse. Statistical significance was determined by Kruskal-Wallis with Conover-Iman multiple comparisons procedure using Benjamini-Hochberg for controlling false discovery rate (panels B and D), two-way repeated-measures ANOVA with Tukey’s multiple-comparisons test (panel F), and the log-rank (Mantel-Cox) test (panel G). *P < 0.05, **P < 0.01, ***P < 0.001, ****P < 0.0001; ns, not significant.

We then challenged mice with WT HSV-2 (strain 186 syn^+^) four weeks after BEACON delivery (**Fig. 3E**). BEACON-treated animals exhibited markedly lower pathology scores and enhanced survival relative to saline, free CpG ODN, or CXCL9-treated controls (**Fig. 3F-G**). By contrast, formulations using CXCL10 (which does not form stable nanoparticles) failed to confer protection against WT HSV-2 (**fig. S4F-H, fig. S2E-G**). Collectively, these results indicate that optimal nanoparticle formulation is critical for effective mucosal delivery, selective immune cell targeting, and establishing protective CD8^+^ T_RM_ populations.

### Vaginal mucosal boosting with HSV glycoproteins and BEACON elicits robust HSV-2 protection

Although “prime and pull” can recruit HSV-specific CD8^+^ T cells and improve disease outcomes, it has no effect on viral titers in vaginal secretions or on local antigen-specific antibody levels in the vaginal mucosa (*16*). We next asked whether co-administering BEACON with HSV glycoprotein antigens could improve vaccine-mediated protection (*25*). Using HSV-2 glycoprotein D (gD), currently in clinical trials (*43*, *44*), we evaluated the efficacy of vaginal boosting with BEACON compared to traditional intramuscular boosting. Following intramuscular priming with gD mRNA-LNPs, WT mice received one of the following boosts: (i) intramuscular gD mRNA-LNPs; (ii) intravaginal gD protein alone, (iii) intravaginal BEACON alone; or (iv) intravaginal gD protein co-administered with BEACON (**Fig. 4A**).

**Figure 4.**
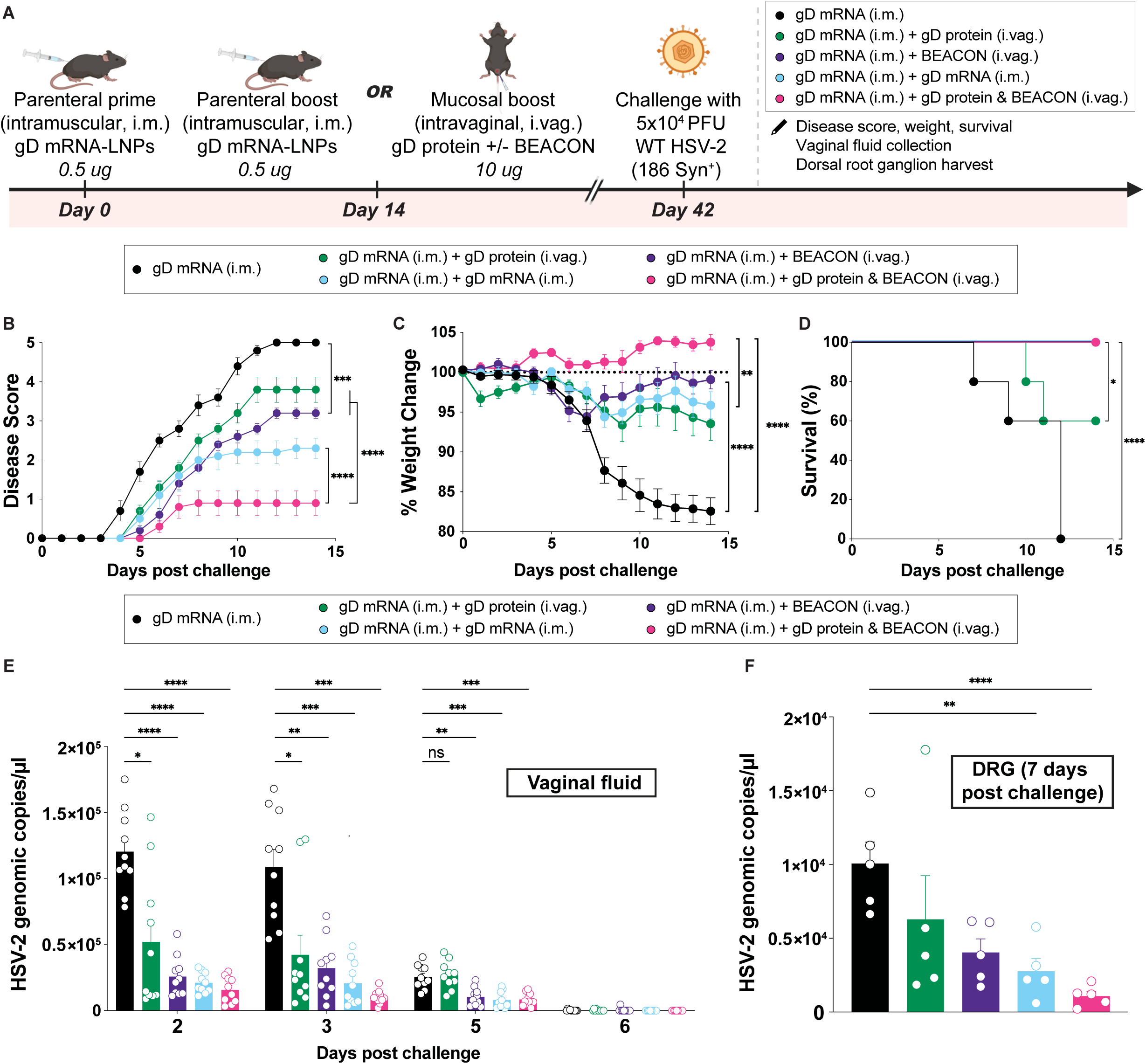
Intravaginal boosting with HSV glycoprotein antigen and BEACON significantly enhances protection against lethal genital HSV-2 challenge compared to parenteral boosting. **(A)** Experimental schematic. Naïve C57BL/6 mice (n = 10 per group) were primed intramuscularly (i.m.) with gD2 mRNA-LNPs (0.5 µg) and boosted with saline, gD2 mRNA-LNPs (0.5 µg, i.m.), gD2 protein (10 µg, i.vag.), BEACON (10 µg, i.vag.), or gD2 protein and BEACON (10 µg, i.vag.). Four weeks post boost, mice were challenged with 5 × 10^4^ PFU of HSV-2 (strain 186 syn^+^). **(B)** Clinical disease scores, **(C)** percent weight change over time and **(D)** Kaplan-Meier survival curves following WT HSV-2 challenge. **(E)** Genomic copies of HSV-2 titers from vaginal washes collected at the indicated time points post challenge with HSV-2. **(F)** HSV-2 DNA in DRG samples collected on day 7 post-challenge from a separate, parallel cohort of mice (n=5 per group) euthanized specifically for DRG viral DNA quantification. In all bar graphs, each point represents the results from one mouse. Data are shown as mean ± SEM. Disease scores (B) and weight loss curves (C), and viral genomic copies (E) were analyzed by two-way repeated-measures ANOVA with Tukey’s multiple-comparisons test. DRG viral DNA was analyzed by Kruskal-Wallis with Conover-Iman multiple comparisons procedure using Benjamini-Hochberg for controlling false discovery rate (panel F). Survival differences were assessed by the log-rank (Mantel-Cox) test (D). *P < 0.05, **P < 0.01, ***P < 0.001, ****P < 0.0001; ns, not significant.

Intravaginal administration of gD protein with BEACON conferred the greatest reduction in disease severity (**Fig. 4B**). None of the mice in the vaginal boosting group displayed overt signs of illness (e.g., ruffled fur, hunched posture, and/or weight loss), whereas 30% of mice in the intramuscular boosted group developed these signs (**Fig. 4C**). By contrast, 40% of mice receiving gD protein alone and 100% of the prime-only controls developed severe disease and required euthanasia (**Fig. 4D**). To assess viral control, we measured HSV-2 DNA in vaginal washes via quantitative PCR (qPCR). Mice boosted vaginally with gD and BEACON exhibited the most rapid and sustained reductions in viral shedding (**Fig. 4E**). Analysis of DRG at day seven post-challenge revealed an ∼80-fold reduction in viral genome copies in the vaginally boosted group relative to controls (**Fig. 4F**).

Furthermore, longitudinal monitoring revealed that 80% of mice in the vaginal boost group survived through six months post-infection without signs of chronic disease or neurological sequelae (*45*) (**Fig. 5A-B**). By contrast, only 40% of mice boosted intramuscularly with gD mRNA-LNPs survived long term (**Fig. 5B**). Mice in the vaginally boosted cohort also exhibited normal weight gain consistent with age-matched, uninfected controls, whereas animals boosted intramuscularly exhibited limited weight recovery (**Fig. 5C**). Mice boosted intramuscularly and euthanized six months post virus challenge exhibited increased inflammation in the vaginal tract with multifocal submucosal neutrophilic infiltrates and increased lymphocytic infiltration in the urinary bladder (**fig. S5A-B**). To determine whether specific inflammatory phenotypes were associated with long-term ganglionic viral burden, we harvested lumbar and thoracic DRG from survivors six months after challenge and quantified HSV-2 genomes by qPCR (**Fig. 5D, fig. S6A-B**). HSV-2 DNA was detected in lumbar DRG in both boosted groups (**Fig. 5D**). HSV-2 genome copies were also detected in thoracic DRG of intramuscularly boosted mice; levels in intravaginally boosted mice were below the limit of detection (**Fig. 5D**). Consistent with these findings, single-molecule fluorescent in situ hybridization (FISH) targeting the HSV-2 latency-associated transcript (LAT) revealed punctate signals in DRG from intramuscularly boosted mice but not from intravaginally boosted or mock-infected controls (**Fig. 5E, fig. S6C**). These results suggested that enhanced early mucosal control limits the establishment of transcriptionally active latency.

**Figure 5.**
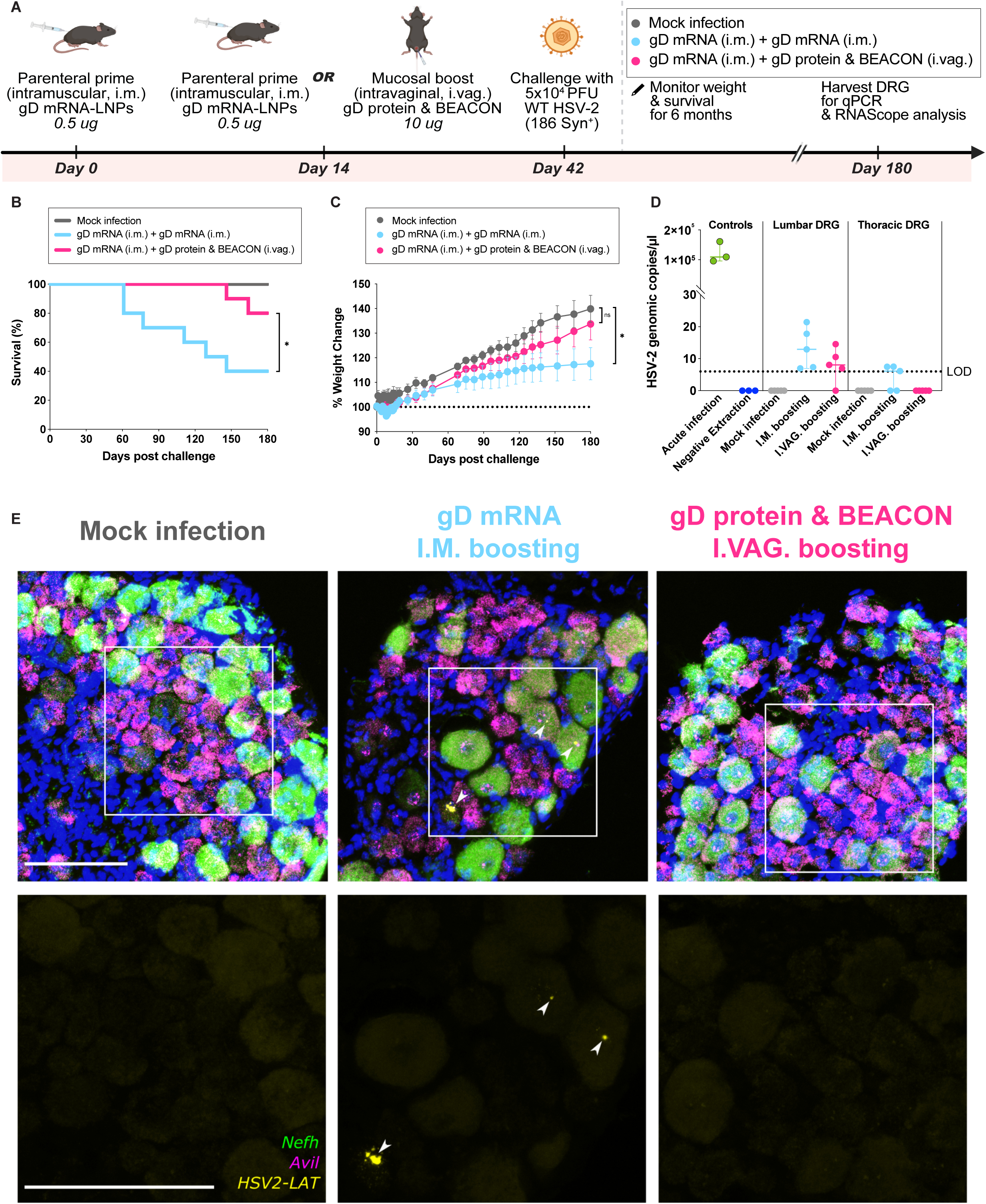
Mucosal boosting is associated with reduced long-term viral detection and LAT signal at six months post HSV-2 challenge. **(A)** Experimental schematic. Naïve C57BL/6 mice (n = 10 per group) were primed intramuscularly (i.m.) with gD2 mRNA-LNPs (0.5 µg) and boosted with gD2 mRNA-LNPs (0.5 µg, i.m.) or gD2 protein and BEACON (10 µg each, i.vag.). Four weeks post boost, mice were challenged with WT HSV-2 and monitored for six months. Surviving mice were euthanized at day 180, and lumbar and thoracic DRG were harvested for HSV-2 genome quantification by qPCR and for spatial analysis by FISH. **(B)** Kaplan-Meier survival curves and **(C)** longitudinal weight monitoring through day 180 post challenge; dotted line indicates baseline weight (100%) at the time of the challenge. **(D)** HSV-2 genomic copies in lumbar and thoracic DRG with the assay limit of detection (LOD) indicated. **(E)** Representative FISH images of lumbar dorsal root ganglia for each group with probes directed against neurofilament heavy chain (*Nefh*; green), advillin (*Avil*; magenta), or the HSV-2 *LAT* (yellow). Images shown in the top row depict the composite overlay of all channels, while those in the bottom row show only the results from the LAT channel, with enlarged regions of interest delimited by boxes in the top row images. Scale bars = 100 µm and apply to all images within the same row. Bars included in the dot plots indicate the mean ± SEM. Survival differences (B) were assessed by the log-rank (Mantel-Cox) test. Weight curves (C) were analyzed by two-way repeated-measures ANOVA. *P < 0.05; ns, not significant.

These findings were corroborated using HSV-2 gB as an alternative immunogen (**fig. S7A**). Vaginal co-delivery of gB protein with BEACON outperformed intramuscular gB mRNA-LNP boosts (**fig. S7B-D**). However, consistent with previous findings (*46*), gD-based immunization provided more robust protection (**Fig. 4B-D**, **fig. S7B-D**). Collectively, these findings show that mucosal boosting with HSV glycoprotein antigens and BEACON provides better protection against HSV-2 challenge than traditional intramuscular methods, with better viral control and improved long-term outcomes.

### Vaginal boosting with BEACON enhances local antibody responses crucial for protection against HSV-2

Mucosal antibody responses play a critical role in preventing HSV-2 infection at the site of viral entry (*18*, *19*). To assess whether vaginal boosting with BEACON enhances humoral immunity, we immunized mice with the same regimens as described in **Fig. 6A**; gD-specific IgG and IgA titers in serum and vaginal fluid were determined by ELISA. While intramuscular boosting with gD mRNA-LNPs resulted in the highest gD-specific serum IgG and IgA titers (**Fig. 6B**), intravaginal boosting with gD protein and BEACON led to markedly elevated levels of antigen-specific IgG and IgA in the vaginal mucosa (**Fig. 6C**).

**Figure 6.**
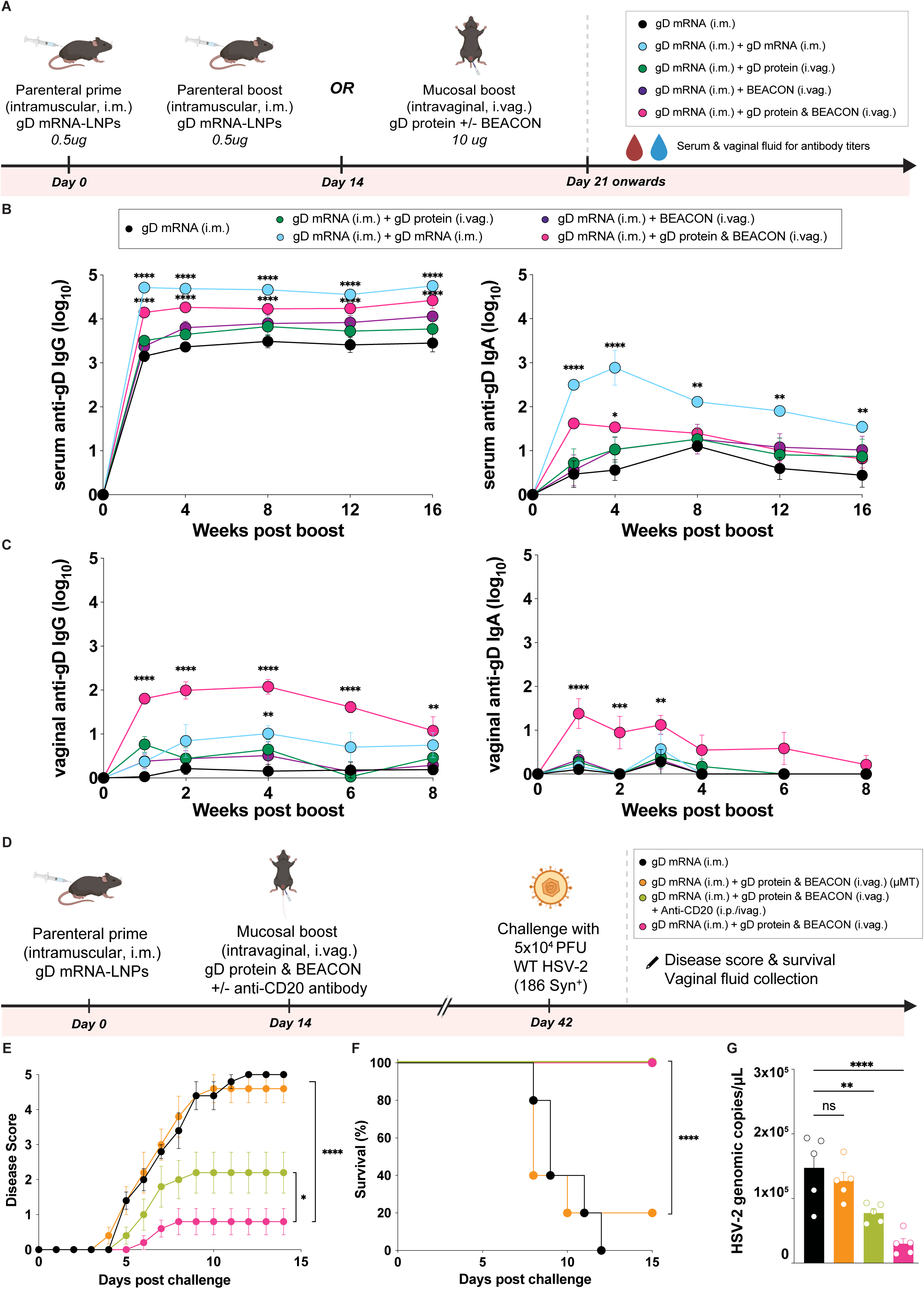
Local antigen availability upon mucosal boosting promotes durable humoral immune responses in the vaginal tract that are indispensable for protection against WT HSV-2 challenge. **(A)** Experimental schematic. Naïve C57BL/6 mice (n = 5 per group) were primed intramuscularly (i.m.) with gD2 mRNA-LNPs (0.5 µg) and boosted with saline, gD2 mRNA-LNPs (0.5 µg, i.m.), gD2 protein (10 µg, i.vag.), BEACON (10 µg, i.vag.), or gD2 protein and BEACON (10 µg, i.vag.) in combination. Serum was collected at 2, 4, 8, 12, and 16 weeks and vaginal fluid was collected at 1, 2, 4, 6, and 8 weeks post-boosting. gD-specific IgG and IgA titers in **(B)** serum and **(C)** vaginal fluid were measured by ELISA at the indicated timepoints. **(D)** Experimental schematic for B cell depletion studies. WT or μMT mice were treated with anti-CD20 antibody or isotype control prior to intravaginal boosting. **(E)** Clinical disease scores and **(F)** survival of mice described in (D) following HSV-2 challenge. Data are representative of two independent experiments (n = 5 mice per group). **(G)** HSV-2 genome copies in vaginal washes collected on day 2 post-challenge. Each point in the bar graphs represents one mouse; bars indicate mean ± SEM. For time-course plots, points indicate mean ± SEM. Antibody kinetics were analyzed by two-way repeated-measures ANOVA with Dunnett’s multiple-comparisons test (B, C). Disease scores were analyzed by two-way repeated-measures ANOVA with Tukey’s multiple-comparisons test (E). Survival differences were assessed by the log-rank (Mantel-Cox) test (F). Quantification of vaginal viral genomes was analyzed by Kruskal-Wallis with Conover-Iman multiple comparisons procedure using Benjamini-Hochberg for controlling false discovery rate (G). *P < 0.05, **P < 0.01, ***P < 0.001, ****P < 0.0001; ns, not significant.

We generated gD tetramers and quantified gD-specific germinal center (GC) B-cells in the draining iliac lymph nodes (LNs) (**fig. S8A**). Intravaginal administration of gD protein and BEACON increased the number of gD-specific GC B-cells relative to intramuscular boosting or prime-only controls (**fig. S8B-C**). Because HSV-2 gD-specific B cells were rarely found in the female reproductive tract (FRT), which limits direct quantification in vaginal tissue, we used a surrogate antigen to test whether mucosal boosting could enhance local antigen-specific B cell accumulation. Hen egg lysozyme (HEL)-specific B cells from MD4 transgenic mice were adoptively transferred into WT recipients before vaccination (intramuscular HEL mRNA priming followed by intravaginal HEL protein and BEACON boost). Antigen-specific B-cells were then enumerated in the FRT (**fig. S8D**). We observed a trend toward increased HEL-specific B-cells in the FRT after mucosal boosting (**fig. S8E-F**).

We next assessed the contribution of B cells using both an antibody-mediated depletion strategy and B cell-deficient μMT mice. WT and μMT mice were primed intramuscularly with gD mRNA and boosted intravaginally with gD protein and BEACON; WT mice received anti-CD20 antibody or isotype control before intravaginal boosting (**Fig. 6D**, **fig. S9A**). Despite residual B cells, anti-CD20-treated mice exhibited significantly elevated disease scores (**Fig. 6E**) and viral loads (**Fig. 6G**), but no change in survival (**Fig. 6F**). Although gD-specific IgG and IgA titers in the vaginal fluid were reduced following anti-CD20 administration, responses remained higher than those observed in mice that received only intramuscular priming (**fig. S9B**). By contrast, no gD protein/BEACON-mediated protection was observed in intravaginally boosted μMT mice over priming-only controls (**Fig. 6E-G**). Together, these data indicate that B cells are required for effective mucosal protection.

Finally, to determine whether this effect applies to other antigens, we also evaluated mucosal antibody responses following gB-based immunization (**fig. S10A**). Similar to the gD findings, mice primed intramuscularly with gB mRNA-LNPs and boosted intravaginally with gB protein and BEACON developed significantly elevated levels of gB-specific antibodies in the vaginal fluid, while systemic titers remained unchanged (**fig. S10B-C**).

### Vaginal boosting with BEACON increases CD8^+^ T_RM_ cells and supports viral protection

To test the importance of CD8^+^ T_RM_ cells in BEACON-mediated protection, we adoptively transferred congenically-marked gBT-I CD8^+^ T cells into naïve mice, followed by intramuscular priming with gB mRNA-LNPs. Mice then received one of four boosts: (i) intramuscular gB mRNA-LNPs; (ii) intravaginal gB protein alone; (iii) intravaginal BEACON alone; or (iv) intravaginal co-delivery of gB protein with BEACON (**Fig. 7A**).

**Figure 7.**
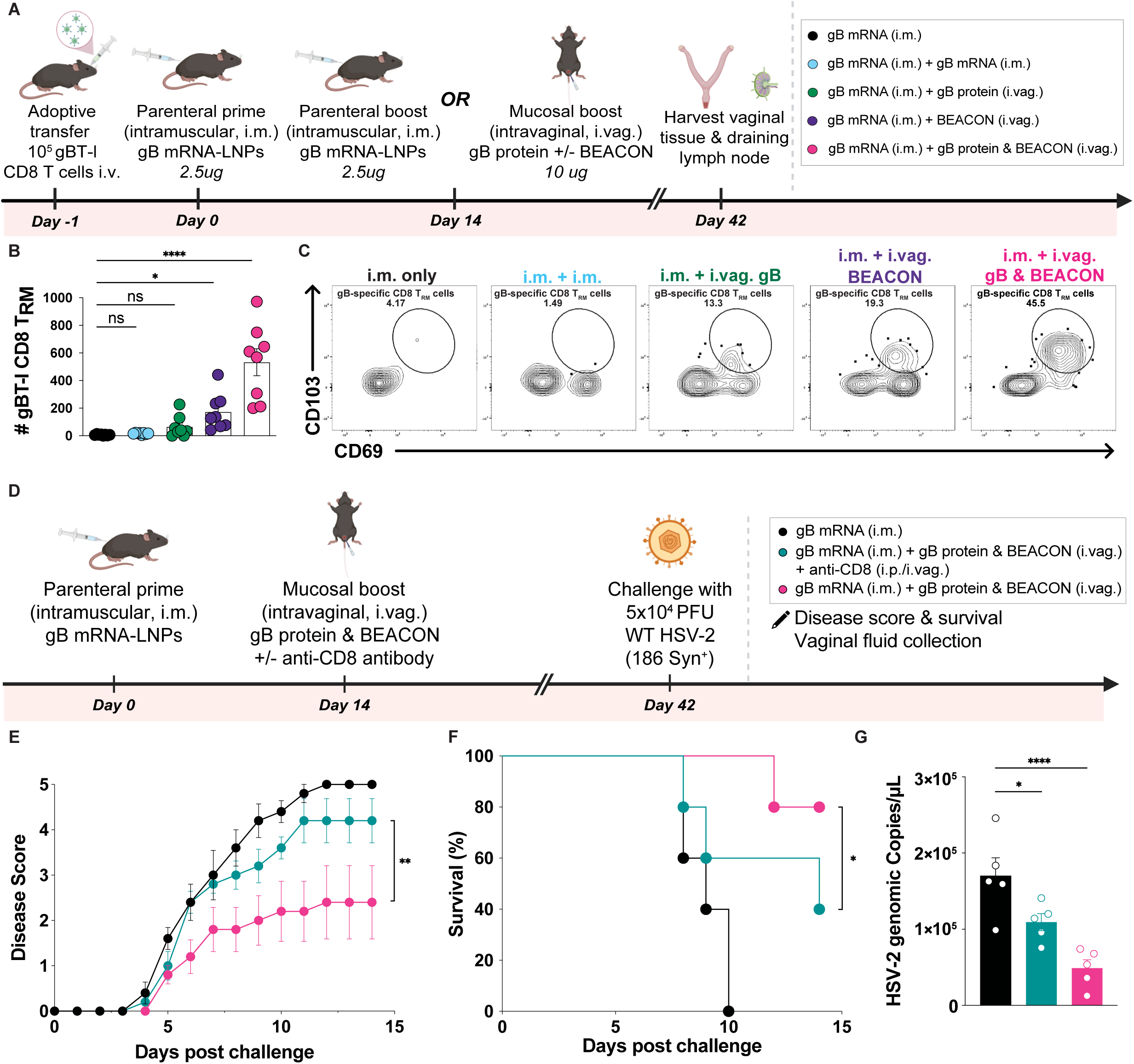
Durable CD8^+^ T_RM_ cell populations generated by mucosal boosting with BEACON contribute to vaccine-mediated protection against WT HSV-2 viral challenge. **(A)** Experimental schematic. Naïve C57BL/6 mice (n = 8 per group) were primed intramuscularly (i.m.) with gB2 mRNA-LNPs (2.5 µg) and boosted with saline, gB2 mRNA-LNPs (2.5 µg, i.m.), gB2 protein (10 µg, i.vag.), BEACON (10 µg, i.vag.), or gB2 protein and BEACON (10 µg, i.vag.) in combination. Four weeks post boost, **(B)** the number of CD8**^+^**T_RM_ (CD69^+^ CD103^+^ gBT-I) cells was quantified in the vaginal tract, with **(C)** representative flow cytometry plots gated on gBT-I tetramer^+^ CD8^+^ T cells shown. Numbers are percent CD69^+^ and CD103^+^ cells. **(D)** Experimental schematic. Naïve C57BL/6 mice (n = 5 per group) underwent priming immunization intramuscularly with gB2 mRNA-LNPs (2.5 µg) followed by intravaginal boosting with gB2 protein and BEACON (10 µg) with or without anti-CD8 antibody treatment. Four weeks post boost, mice were challenged with WT HSV-2. **(E)** Disease scores and **(F)** Kaplan-Meier survival curves in mice following WT HSV-2 challenge. **(G)** HSV-2 genome copies in vaginal washes collected on day two post-challenge. Data in (E-F) are representative of two independent experiments (n = 5 mice per group). For bar graphs, each point represents one mouse and bars indicate mean ± SEM. CD8**^+^** T_RM_ quantification and vaginal viral genomes were analyzed by Kruskal-Wallis with Conover-Iman multiple comparisons procedure using Benjamini-Hochberg for controlling false discovery rate (B, G). Disease scores were analyzed by two-way repeated-measures ANOVA with Tukey’s multiple-comparisons test (E). Survival differences were assessed by the log-rank (Mantel-Cox) test (F). *P < 0.05, **P < 0.01, ***P < 0.001, ****P < 0.0001; ns, not significant.

Vaginal tissues were analyzed by flow cytometry at four weeks post-boost. Mice receiving vaginal gB protein and BEACON exhibited a three- to four-fold increase in gBT-I CD8^+^ T_RM_ cells compared to mice treated with BEACON alone (**Fig. 7B**). These gBT-I CD8^+^ T cells displayed a canonical T_RM_ phenotype (CD69^+^ CD103^+^) and were excluded from intravascular anti-CD45 labeling (**Fig. 7C**). Notably, none of the other vaccination regimens, including intramuscular boosting, resulted in CD8^+^ T_RM_ populations above baseline levels. The increase in T_RM_ cells was localized to the vaginal mucosa; the number of gBT-I CD8^+^ T cells in draining iliac lymph nodes (LNs) remained unchanged across all four groups (**fig. S11A-C**). CD4^+^ T cell frequencies were also unaltered in both the vaginal and LN compartments (**fig. S11B-D**). To assess the durability of the vaginal CD8^+^ T_RM_ response elicited by mucosal boosting, we analyzed tissues on day 98 after intravaginal boosting with gB protein and BEACON. Intravaginal gB protein and BEACON maintained significantly higher numbers of gBT-I CD8^+^ T_RM_ cells in the vaginal tract compared to priming-only or intramuscularly boosted controls (**fig. S11E-F**). Congenic tracking revealed that both adoptively transferred (CD45.1^+^) and endogenous (CD45.2^+^) gB tetramer^+^ CD8^+^ T cells acquired a CD69^+^CD103^+^ T_RM_ phenotype in the vaginal tract. By contrast, the long-lived vaginal T_RM_ pool was dominated by the adoptively transferred gBT-I population, with a smaller number of endogenous gB-specific T cells, suggesting that mucosal boosting primarily recruits systemically primed effectors (**fig. S11G-H**).

To assess the contribution of CD8^+^ T cells to mucosal vaccine-mediated protection, we treated mice with anti-CD8 depleting or isotype-matched control antibodies prior to HSV-2 challenge (**Fig. 7D, fig. S12A-B**). CD8-depleted mice exhibited significantly increased disease severity and reduced survival compared to isotype-treated controls (**Fig. 7E-F**). Viral genome copies in vaginal washes on day two post-challenge were elevated in CD8-depleted mice compared to isotype-treated controls (**Fig. 7G**). These findings indicate that while CD8^+^ T cells are key effectors of vaccine-induced protection, additional immune mechanisms are needed to promote viral control. Similar results were obtained when gD was used as the immunogen; CD8^+^ T cell depletion led to elevated disease scores but did not have a substantial impact on short-term survival (**fig. S12C-E**). CD8^+^ T cell depletion also reduced vaginal gD-specific IgG levels (**fig. S12F**), suggesting a role for these cells in promoting local humoral responses (*47*). These results underscore the essential but non-exclusive role of CD8^+^ T cells in mediating protection following mucosal boosting.

These findings show that local CD8^+^ T_RM_ and antibody responses both contribute to BEACON-induced protection, and emphasize the importance of engaging multiple immune arms at mucosal surfaces against genital herpes.

## DISCUSSION

In this study, we show that combining parenteral priming with localized vaginal boosting using BEACON with HSV glycoproteins yields significant protection against genital HSV-2 infection. This protection is associated with increased numbers of vaginal CD8 ^+^ T_RM_ cells and elevated mucosal antibody responses, highlighting the importance of local immune effectors for protective antiviral defense (*20*, *48*).

We designed BEACON by leveraging electrostatic interactions between CpG ODN and the chemokine CXCL9. These nanoparticles exhibited stable morphology and function, enabling effective uptake, TLR9-mediated activation of APCs, and chemokine-directed recruitment of effector lymphocytes. When administered intravaginally, BEACON was preferentially taken up by DCs and monocytes, resulting in significantly reduced inflammation compared to free CpG ODN. This altered biodistribution likely contributes to the improved tolerability and immune activation observed in vivo (*21*). Notably, the cationic C-terminal region of CXCL9 was required for nanoparticle formation; BEACON generated using CXCL10 or truncated CXCL9 failed to form stable particles and did not confer protection. These data emphasize the structural features of the chemokine component that directly influence particle formation and immunological function.

Following intramuscular priming, intravaginal boosting with BEACON and viral glycoprotein antigen induced a substantial increase in HSV-specific CD8^+^ T_RM_ cells in the vaginal tissue (*16*, *49*). This response was not induced by CXCL9 or CpG ODN alone, nor was it observed after intramuscular boosting, indicating that both local antigen and adjuvant signals are required to establish durable CD8^+^ T_RM_ populations. Our data support a model in which mucosal boosting primarily leads to recruitment and locally reprogramming of systemically primed effectors rather than driving a second systemic expansion. We determined that the increase in T_RM_ cells was localized to the vaginal mucosa; no corresponding increases in gBT-I CD8^+^ T cell numbers were observed in draining iliac LNs. Congenic tracking revealed that the long-lived gB tetramer^+^ vaginal T_RM_ pool was dominated by the donor-derived population primed intramuscularly. Moreover, the vaginal T_RM_ population persisted for ∼100 days after boosting, indicating that mucosal boosting can durably seed the genital tract with resident effectors. Although recombinant protein is often ineffective at eliciting CD8 responses in these settings, we hypothesize that BEACON enhances local uptake and activation of APCs and promotes cross-presentation of soluble glycoprotein by specialized DC subsets in the draining LNs and/or the vaginal mucosa, thereby providing the signals necessary to induce CD69^+^ and CD103^+^ residency programs in T cells. This hypothesis is consistent with prior work showing that HSV antigens are cross-presented preferentially by CD301b^+^ DCs and that T_RM_ restimulation in the FRT requires MHC-I expression on CD301b^+^ DCs (*50*). It remains an open question whether the impact of the BEACON booster depends solely on recruiting pre-formed T cells into the FRT or on expanding them systemically or within the FRT. Resolving this issue will require temporal and cell-tracking approaches.

We also observed strong mucosal antibody responses following vaginal boosting. Intravaginal delivery of gD protein with BEACON produced markedly elevated levels of antigen-specific IgG and IgA in the vaginal mucosa over time, whereas systemic antibody titers were highest after intramuscular mRNA-LNP boosting. This compartmentalization suggests that direct antigen exposure in the mucosa drives robust luminal antibody responses and supports a model in which mucosal boosting selectively enhances barrier antibodies without proportionately altering systemic responses (*19*, *20*). Mucosal boosting increased gD-specific GC B cells in the vagina-draining LNs. While direct enumeration of HSV gD-specific B cells in the FRT was limited by low recovery, preliminary results from surrogate adoptive transfer experiments suggested that mucosal boosting can promote antigen-specific B cell accumulation in the FRT.

BEACON-mediated protection requires coordinated local T and B cell immunity. CD8 ⁺ T cell depletion before challenge increased disease severity and early viral burden, demonstrating that CD8⁺ T cells are critical for limiting early replication in the mucosa (*16*, *49*, *50*). In addition, although mice subjected to partial B cell depletion with an anti-CD20 antibody survived, they exhibited higher disease scores and increased viral shedding. These findings support the existence of an antibody threshold that prevents lethal neurologic disease despite incomplete local control. Strikingly, the protective benefit of intravaginal gD protein plus BEACON was abrogated in B-cell-deficient μMT mice, indicating that B cells and mucosal antibody responses are required for optimal protection in this model. Together, these data support a framework in which mucosal antibodies and resident T cells act in concert: antibodies neutralize viruses at the epithelial surface and limit early spread, while T_RM_ cells rapidly eliminate infected cells to contain breakthrough infections and prevent the establishment of latency within the DRG (*18*).

A comparison of glycoprotein antigens also suggests a role for distinct effector hierarchies in promoting antiviral protection. In WT C57BL/6 mice, gB contains a dominant CD8 epitope, whereas gD elicits comparatively weak CD8 responses. Nonetheless, gD-based mucosal boosting provided particularly strong protection, consistent with a prominent role for neutralizing antibodies and CD4 - dependent helper T cell activity against this glycoprotein necessary for viral entry (*46*). Similarly, the observation that CD8^+^ T cells contribute to disease control even with gD-based regimens supports the idea that mucosal boosting can recruit and retain a broader population of CD8 ^+^ effector cells that become functionally relevant once infection is established.

Longitudinal monitoring revealed additional separation between mucosal and intramuscular boosting strategies that was not predicted by serum titers alone. This divergence in outcome despite comparable serologic profiles challenges the historical reliance on serum neutralizing antibody as a correlate of protection and points to the necessity of evaluating local immune parameters in mucosal HSV vaccine design (*12*). Mucosal boosting reduced the number of viral genome copies detected in DRG early after challenge and was associated with improved long-term weight recovery and survival. Although spontaneous HSV reactivation is not a well-described feature of the mouse vaginal challenge model, we observed signs of transcriptionally active latency and viral spread from local to distal DRG in parenterally immunized but not mucosally boosted mice. Collectively, these results suggest that the ganglionic distribution of viral genomes is broader when barrier immunity at the portal of entry is suboptimal. Notably, no LAT signals were detected in DRG from intravaginally boosted mice; low-level HSV DNA remained detectable in lumbar ganglia, suggesting an abortive latency rather than a fully established, transcriptionally active latent virus reservoir. Limiting early replication at the portal of entry constrains the extent and distribution of ganglionic seeding (*51*). Future studies using ex vivo ganglionic reactivation assays and additional temporal profiling will be required to determine whether mucosal boosting reduces the frequency of reactivation -competent latent virus. Furthermore, additional mechanistic studies will be needed to define the cellular composition and antigen specificity of long-term vaginal and bladder infiltrates and establish causal links to ganglionic viral burden.

Our work has several broader implications. First, it provides a platform for decoupling the immunologic requirements for priming and boosting. Our results reveal that systemic priming can generate a broad effector pool and that mucosal boosting can localize and amplify relevant effectors at the site of infection (*34*). Second, our findings demonstrate that the reactogenicity of CpG ODN can be fine-tuned through nanoparticle formulation to mitigate inflammation without sacrificing efficacy, a critical consideration in clinical translation (*21*, *22*). Third, our findings offer a generalizable strategy for targeting other sexually transmitted pathogens, including HIV-1, human papillomavirus, and chlamydia, in which mucosal immunity is essential but poorly induced by parenteral vaccines (*35*).

Clinical translation of this approach will require careful consideration of safety and biological heterogeneity. Despite the availability of intravaginal hormones, contraceptives, and anti-infective products, there are currently no intravaginal vaccines or immunostimulants licensed for human use. Furthermore, results from prior studies on topical interventions have shown that mucosal inflammation can undermine their efficacy (*52–54*). Accordingly, early-phase clinical development of a cervicovaginal booster would need to prioritize local tolerability and mucosal safety endpoints, with documentation of genital adverse events. Efficacy and tolerability may also vary with menstrual cycle stage, post-menopausal status, hysterectomy, pregnancy, microbiome heterogeneity, and intrauterine device use, all of which can shape baseline inflammation and barrier function. In addition, species-specific differences in TLR9 expression and CpG motif preferences will require further validation (*55*). To facilitate translation, our platform has also been designed using a clinically approved, human-active TLR9 agonist (CpG 1018) and human CXCL9. Furthermore, although dosing and formulation will require further optimization, we anticipate that the delivery-related advantages of BEACON will be preserved in human cervicovaginal mucosa. Finally, whether BEACON may be used for therapeutic vaccines requires future research (*56*). This study has several limitations. Although we demonstrate the durability of vaginal CD8^+^ T_RM_ cell populations and follow animals for six months after challenge, longer-term persistence and recall under conditions that closely mimic human physiology, including hormonal cycling, diverse microbiota, and repeated exposure, remain to be determined. Furthermore, while our long-term observational study suggests that mucosal boosting can suppress reactivation-competent virus within the DRG, we did not formally test the impact of this strategy on vaginal viral shedding (*2*). Testing our strategy in guinea pigs, which exhibit recurrent HSV-2 disease, would provide important insight into this question (*56*). Finally, methods that might be used to adapt this strategy for self -administered or clinically deployable formulations, including slow-release platforms or thermally stable vaccine suppositories, require further development (*25*, *28*, *57*).

In conclusion, our study demonstrates that localized mucosal boosting with HSV glycoprotein antigens and BEACON following systemic priming elicits potent tissue-resident T cell and antibody responses that are essential for protection against genital HSV-2 infection. By revealing mechanistic requirements for durable barrier immunity, these results suggest that local antigen delivery in the context of a well-tolerated mucosal adjuvant can overcome key limitations of current parenteral HSV vaccine strategies.

## MATERIALS & METHODS

### Study design

The primary aim of this research was to assess the impact of vaginal boost compared to traditional intramuscular boost immunizations on protection in mice challenged with a lethal dose of WT HSV-2. To achieve this goal, we developed a nanoparticle mucosal adjuvant, performed parenteral prime-mucosal boost immunization experiments with clinically relevant immunogens, and evaluated cellular and humoral responses over time. Our mechanistic studies examined the protective role of local antigen-specific B and T cell responses using cell-depleting antibodies and transgenic mice.

### Immunogens

Messenger RNAs encoding HSV-2 glycoproteins gD2, gB2, and HEL (for surrogate antigen studies) were purchased from Cellerna Bioscience (Baesweiler, Germany) and encapsulated in LNPs. Additional details are provided in the Supplementary Materials.

The secreted ectodomain of recombinant HSV-2 gD2 was expressed in HD Chinese Hamster Ovary (CHO)-S cells (GenScript, Piscataway, NJ, USA). The mature gD2 ectodomain spans residues 26-310 (UniProt P03172) and includes a C-terminal histidine (His)_6_ tag; R52Q is the sole polymorphism. Expression was achieved by transient transfection and culture of CHO cells in suspension with the codon-optimized (His)₆-tagged gD2 ectodomain gene fragment subcloned into pcDNA3.1(+). Supernatants were harvested on day 10, and protein was purified by nickel-nitrilotriacetic acid (Ni-NTA) affinity chromatography followed by buffer exchange into phosphate-buffered saline (PBS). Purity (>95%) was confirmed by SDS-PAGE and Western blotting. Protein concentration was determined by absorbance at A_280_. HSV-2 gB2 protein expressed in and purified from HEK293 (ACROBiosystems, Newark, DE, USA; catalog GLB-H52H3) includes the full ectodomain of HSV-2 strain 333 gB (Uniprot P06484) with a C-terminal His_6_ tag. Monodispersity and purity were verified by multi-angle light scattering.

### Nanoparticle adjuvant preparation

Recombinant murine CXCL9 and CXCL10 (PeproTech) were reconstituted at 1 mg/mL in 25 mM HEPES (pH 7.4). Stock solutions of CpG-ODN 1826 (Invivogen) were prepared at 1 mM in endotoxin-free water. Equal volumes of chemokine and CpG ODN were mixed at the indicated molar ratios. Nanoparticle morphology was determined, and the solution was rendered isotonic by the addition of dextrose (10% w/v).

### Mice

All animal studies were conducted in accordance with National Institutes of Health (NIH) guidelines and approved by the Yale Institutional Animal Care and Use Committee. Eight-to-12-week-old female C57BL/6NCrl (WT) mice were purchased from Charles River. Mice were housed under specific pathogen-free conditions with ad libitum access to food and water and maintained on a 12 -hour light/dark cycle. CD45.1^+^ gBT-I TCR transgenic mice were provided by F. R. Carbone and W. R. Heath and bred in-house on a C57BL/6 background. MD4 BCR transgenic mice (strain 002595, Jackson Laboratory) were used as donors for HEL-specific B-cell adoptive transfer experiments as indicated.

### Immunizations and sample collections

Mice were anesthetized with isoflurane. To synchronize estrous cycles, mice received subcutaneous injections of 2 mg Depo-Provera (medroxyprogesterone acetate; Prasco, Mason, OH) diluted to 20 mg/mL in sterile PBS, five days before vaginal immunization or HSV-2 challenge. For systemic priming, mice were immunized intramuscularly (quadriceps) with gD2 (0.5 μg) or gB2 (2.5 μg) mRNA-LNPs (Cellerna Bioscience) in 50 μL PBS. For mucosal boosting, 10 μg recombinant HSV-2 glycoprotein (GenScript) in 20 μL PBS was administered intravaginally alone or in combination with BEACON. When co-administered, HSV protein and BEACON were mixed gently for 30 min at room temperature. Mice were held inverted for approximately 1 min post-administration to facilitate complete absorption. For surrogate antigen experiments, recipient mice were primed intramuscularly (2 μg each) with HEL mRNA-LNPs (Cellerna Bioscience) in 50 μL and boosted on day 14 either intramuscularly with HEL mRNA-LNPs or intravaginally with HEL protein and BEACON (10 μg). Blood was collected retro-orbitally under anesthesia. Sera were isolated by centrifugation and stored at −80°C. Vaginal fluid was collected using Puritan® polyester swabs rotated within the vaginal tract and subsequently eluted with 50 μL PBS, and stored at −80°C. Dorsal root ganglia (DRG) were collected on day 7 post-challenge and stored in PBS at −80°C. Iliac and sacral LNs were harvested and treated with protease inhibitor buffer (protease inhibitor cocktail and ethylenediaminetetraacetic acid in PBS with 2% fetal bovine serum [FBS]), processed using a handheld motorized pestle (VWR), and centrifuged at 16,000 g for 5 min at 4°C.

### Flow cytometry

Single-cell suspensions were prepared by digesting minced vaginal tissues in 0.5 mg/mL Dispase II (Roche, Indianapolis, IN) in PBS for 15 min at 37°C followed by 1 mg/mL collagenase D and 30 μg/mL DNase I (Roche) in complete Dulbecco’s Modified Eagle’s Medium (DMEM) for 45 min at 37°C. Digests were filtered through 70 μm strainers (Falcon), washed twice with PBS, and resuspended in fluorescence-activated cell sorting (FACS) buffer (PBS with 5% FBS and 0.09% sodium azide). To assess tissue residency, mice received retro-orbital injections of fluorescein isothiocyanate (FITC)-labeled anti-mouse CD45 (2 µg in 100 µL PBS) three min before tissue harvest; IV label-positive events were excluded from T_RM_ quantification. Cells to be evaluated by flow cytometry were incubated with a fixable Live/Dead viability dye (Zombie Aqua, BioLegend, San Diego, CA, USA) for 10 min at room temperature in PBS, washed twice, and then stained for 45 min at 4°C in FACS buffer with surface antibody panels prepared in Fc-blocking antibody (anti-CD16/32, clone 93, BioLegend; 1:200). To target antigen-specific GC B-cells, antibody-stained cells were distributed evenly, diluted 1:100 in 50 μL of FACS buffer, and exposed to biotinylated gD protein preincubated with BV421-(BioLegend, 405226) or allophycocyanin-(BioLegend 405207) conjugated streptavidin (1:4 molar ratio, 30 min, 25°C). All fluorochrome-conjugated antibodies and clones are listed in the Supplementary Materials. Cutoffs for positive-staining populations were established using isotype controls during initial panel validation. Fluorescence minus one (FMO) controls were used to define gating boundaries in experimental samples. Compensation was performed using single-stained UltraComp eBeads (ThermoFisher Scientific) and verified with FMOs. For quantitative normalization, entire vaginal tracts from individual mice were processed under identical digestion and staining conditions. CountBright absolute counting beads (Invitrogen) were added to each sample before data acquisition to normalize immune cell counts to total tissue equivalents, thereby enabling comparison of absolute cell numbers across treatment groups. Samples were acquired on a BD FACSymphony A5 and analyzed using FlowJo v10 (Treestar, Ashland, OR, USA).

### Antibodies

Antibodies used for flow cytometry are listed in the Supplementary Materials. H-2Kb-gB498-505 tetramers were obtained from the NIH Tetramer Core Facility. Anti-mouse CD8β (Ly-3.2; clone 53-5.8) and anti-mouse CD20 (clone SA271G2) were used for CD8^+^ T cell and B cell depletion studies, respectively. All reagents were used at manufacturer-recommended concentrations.

### Adoptive transfers, viral challenge, and quantification of viral titers, weight, and disease scores

Single-cell suspensions were generated from the spleens of CD45.1^+^ gBT-I donor mice. CD8^+^ T cells were purified by negative selection (STEMCELL Technologies). A total of 1×10^5^ purified gBT-I CD8^+^ T cells were inoculated retro-orbitally into Depo-Provera-treated CD45.2^+^ recipient C57BL/6 mice one day before priming. For HEL-specific B cell adoptive transfer experiments, spleens from MD4 donor mice were processed into single-cell suspensions, and 10^6^ HEL-specific B cells were transferred intravenously into CD45.2 ^+^ recipient mice one day before immunization. For prime and pull experiments, mice were injected subcutaneously with 10^6^ plaque-forming units (PFU) of TK-deficient HSV-2 (strain 186TKΔkpn). For challenge experiments, mice were infected intravaginally with 5×10^4^ PFU of WT HSV-2 (strain 186 syn⁺) in 10 μL PBS. This inoculum corresponds to approximately five times the median lethal dose for this stock (∼1×10^4^ PFU in Depo-Provera-treated female C57BL/6 mice aged 8-12 weeks). All viral stocks were aliquoted and stored at −80°C and were thawed only once to maintain consistency. The titer of the WT HSV-2 challenge stock was determined by standard plaque assay using Vero cells (ATCC CCL-81). Briefly, Vero cells were seeded at 10^5^ per well in 24-well plates and grown to 95% confluence in DMEM supplemented with 10% FBS and 1% sodium pyruvate. Ten-fold serial dilutions of viral stock were prepared in serum-free DMEM, and 100 μL of each dilution was added to duplicate wells following a single PBS wash. Plates were incubated for 1 h at 37°C with gentle rocking (60 rpm), after which the inoculum was aspirated and replaced with 1 mL of 1.2% methylcellulose (MCC) overlay medium (1:1 mixture of 2.4% MCC and DMEM without supplements). Infected cells were incubated for 72 h at 37°C, fixed with 4% paraformaldehyde for 15 min, washed twice with PBS, and stained with 0.5% crystal violet for 15 min at room temperature. Plates were washed with distilled water, and plaques were enumerated manually in wells containing 10-100 discrete plaques. The HSV-2 viral titer (2×10^8^ PFU/mL) was calculated as the (number of plaques × dilution factor)/inoculum volume [mL]. Virus-challenged mice were weighed and monitored daily for 15 days post-infection (and once per week long-term) for survival and evidence of clinical disease. Disease severity was scored using a five-point clinical scale: 0, no signs; 1, mild genital erythema and edema; 2, moderate inflammation; 3, extensive inflammation with grooming defects; 4, hind-limb paralysis; and 5, premoribund. Mice reaching a score of 5 were euthanized for humane reasons. Body weights were recorded daily, and survival data were plotted using Kaplan-Meier curves.

### Quantification of HSV-2 genomes by qPCR

TaqMan qPCR targeting HSV-2 gB was performed in a total volume of 20 µL, using Luna Universal Probe qPCR Master Mix (New England Biolabs, Ipswich, MA, US) with 5 µL template; forward primer (5′-AGAGGGACATCCAGGACTTTG-3′); reverse primer (5′-GAGCTTGTAATACACCGTCAGG-3′); and probe (Cy5)-CCGCCGAACTGAGCAGACACCCGCGC-(IABkFQ) (Integrated DNA Technologies, Coralville, IA, US) (*58*). DNA extraction and amplification details are provided in the Supplementary Materials.

#### ELISA

High-binding 96-well plates were coated overnight at 4°C with 2 μg/mL of gD2 or gB2 in PBS (50 μL/well). Plates were washed, blocked with 2% bovine serum albumin in PBS (200 μL/well, two hours, room temperature, and incubated with serially diluted serum or vaginal fluid samples for two hours. After five washes, plates were incubated with anti-mouse IgG-horseradish peroxidase (HRP) or IgA-HRP (Bio-Rad) at a 1:3000 dilution in blocking buffer for one hour. Plates were developed with a 3,3′,5,5′-tetramethylbenzidine substrate (50 μL/well) and stopped with 35 μL of 2N H_2_SO_4_. Absorbance was read at 450 nm. Endpoint titers were defined as the highest dilution yielding a signal that exceeded two standard deviations above the background level.

#### Fluorescence in situ hybridization (FISH)

Lumbar DRG were dissected from mice into Optimal Cutting Temperature compound (Sakura Finetek USA, Torrance, CA, US) and flash frozen. Frozen sections were cut to 15 µm in a cryostat and mounted onto Superfrost Plus glass slides (ThermoFisher). FISH for mRNAs encoding neurofilament heavy chain (*Nefh*), advillin (*Avil*), or *HSV-2 LAT* was performed with an RNAscope V2 detection kit and probe sets available from the manufacturer (Biotechne, Minneapolis, MN, US). Briefly, cryosections were fixed in 4% paraformaldehyde in PBS for 1 h at 4°C, incubated for 10 min in peroxide, digested with Protease III for 25 min at room temperature, and hybridized with probes for 2 hours at 40°C. Signals were visualized with Opal 520, Opal 570, and Opal 690 dyes according to the manufacturer’s instructions. Images were collected with a 40× oil-immersion objective on a Stellaris 8 confocal microscope (Leica) using identical instrument settings across all specimens. Positive *HSV-LAT* signals were required to be at least 0.8 µm in diameter with voxels of at least 20,000 arbitrary fluorescence units in 16-bit images.

#### Statistical analysis

Statistical analysis and graphing of the experimental data were performed with GraphPad Prism and R. Two-tailed Student’s t tests were used to compare two experimental groups. For more than two groups, Kruskal-Wallis tests using the Conover-Iman multiple comparisons procedure with Benjamini-Hochberg to control the false discovery rate were used to perform multiple comparisons. Two-way repeated-measures ANOVA with Tukey (pair-wise) or Dunnett’s (against control) multiple comparisons tests were used for time series data. Survival differences were determined from Kaplan-Meier plots with the log-rank (Mantel-Cox) test. A value of *P*<0.05 was considered statistically significant.

## Acknowledgements

We thank M. Linehan, H. Dong, P. Lu, S. Tabachnikova, M. Dresler, and P. Baevova for technical support. We would also like to thank V. Davé, M. Grun, and M. Saltzman for helpful discussions. We also acknowledge the Center for Cellular and Molecular Imaging, Electron Microscopy Facility at the Yale School of Medicine, specifically X. Liu and K. Gibson, for assistance with the electron microscopy work.

## Author contributions

S.H.B and A.I conceived the experiments and wrote the manuscript and all authors discussed and commented on it. S.H.B, S.E, I.S.P, N.C, D.K, C.K., C.A.B, C.L, K.S, C.M, P.G, S.L, A.O, A.H, S.F, R.B.O, and W.B.H performed all the experimental work and analyzed the data. C.J.B performed all the pathology work. S.H.B generated the figures and all authors provided feedback on it. A.I acquired funding and supervised the research.

## Data and Materials Availability

All data needed to evaluate the conclusions in the paper are present in the paper and/or the Supplementary Materials.

## Supplementary Materials

Supplementary Text

Figs. S1 to S12

Data file S1

MDAR Reproducibility Checklist

## Supplementary Materials and Methods

### Details of immunogens

Both HSV-2 constructs (gD2 and gB2) encode secreted ectodomains and do not encode the native C-terminal transmembrane or cytoplasmic domains. Messenger RNAs were synthesized directly from the amino-acid sequences based only on codon optimization without any amino-acid substitutions or sequence modifications. No “detoxifying” mutations were introduced in the gD2 herpesvirus entry mediator (HVEM)-binding region, and no pre-fusion stabilizing mutations were introduced into the gB2 sequences. The mRNA sequence corresponding to gD2 (US6) encoded the mature secreted ectodomain of HSV-2 strain 333 gD (GenBank LS480640 and curated as UniProt P03172). The mRNA sequence corresponding to gB2 (UL27) encoded the ectodomain of HSV-2 strain HG52 gB (reference genome GenBank Z86099.2; RefSeq NC_001798). For HEL, the mRNA encoded hen egg lysozyme (P00698) as indicated. Briefly, in vitro transcription of gD2, gB2 and HEL mRNAs was performed using T7 RNA polymerase. The plasmid DNA, containing a T7 promoter, a 5′ untranslated region (5′ UTR), the coding sequence of gB2 or gD2 or HEL (codon-optimized using a proprietary artificial intelligence-based algorithm developed by Cellerna Bioscience), a 3′ UTR, and a poly(A) tail, were utilized as the template for mRNA synthesis. To generate capped transcripts, a Cap 1 analog was incorporated co-transcriptionally. All uridines were fully substituted with N1-methylpseudouridine during transcription to enhance mRNA stability and reduce immunogenicity. A polyadenylated tail was transcribed from the poly(A) sequence encoded within the DNA template. Following transcription, the reaction mixtures were treated with DNase I to degrade the DNA template. Initial purification of the mRNA was performed by lithium chloride precipitation, which selectively precipitates high-molecular-weight RNA while removing residual proteins, enzymes, and free nucleotides. To eliminate any residual double-stranded RNA contaminants, a second purification step was performed using a cellulose-based chromatography column; this resulted in highly pure, single-stranded mRNA suitable for downstream applications. LNPs were formulated via flash nanoprecipitation. Briefly, SM-102 (BroadPharm, San Diego, CA, US), 1,2-distearoyl-sn-glycero-3-phosphocholine (Avanti Polar Lipids, Snaith, UK), cholesterol (Avanti Polar Lipids), and PEG-DMG 2000 (Avanti Polar Lipids) were dissolved in ethanol at a molar ratio of 50:10:38.5:1.5, respectively. The lipid and mRNA solutions were mixed at a nitrogen to phosphorus (N/P) ratio of 6:1 and final RNA concentration of 0.1 mg/mL using an Ignite NanoAssemblr (Precision NanoSystems, Vancouver, Canada) at a flow rate of 12 mL/min and a lipid:RNA flow ratio of 3:1. Formulated LNPs were dialyzed in 8% sucrose and 20 mM Tris-acetate overnight in 3500 MWCO Slide-A-Lyzer dialysis cassettes (ThermoFisher, Waltham, MA, US) and stored at −80°C until use.

### Amino acid (AA) sequences of gD2 and gB2 immunogens

Messenger RNA sequences were synthesized from the AA sequences listed below (codon optimization only). No “detoxifying” substitutions (e.g., in the gD2 herpesvirus entry mediator [HVEM]-binding region) or prefusion-stabilizing mutations (in gB2) were introduced.

### (A) gD2 mRNA-encoded antigen

**gD2 AA sequence (mRNA-encoded ectodomain):**

<u>MGILPSPGMPALLSLVSLLSVLLMGCVAETG</u>KYALADPSLKMADPNRFRGKNLPVLDQLTDPPGVKRV

YHIQPSLEDPFQPPSIPITVYYAVLERACRSVLLHAPSEAPQIVRGASDEARKHTYNLTIAWYRMGDNCAIPITVMEYTECPYNKSLGVCPIRTQPRWSYYDSFSAVSEDNLGFLMHAPAFETAGTYLRLVKINDWTEITQFILEHRARASCKYALPLRIPPAACLTSKAYQQGVTVDSIGMLPRFIPENQRTVALYSLKIAGWHGPKPPYTSTLLPPELSDTTNATQPELVPEDPEDSALLEDPAGTVSSQIPPNWHIPSIQDVAPHHAP

### (B) Recombinant gD2 protein

**Protein expression/purification:** The secreted gD2 ectodomain was produced in CHO-S cells (GenScript). The purified protein corresponds to the mature ectodomain (UniProt P03172 residues Lys26 - Thr310) with a C-terminal His_6_ tag. GenScript identified one polymorphism (R52Q) within the ectodomain; no other substitutions were noted.

### gD2 AA sequence (recombinant protein)

ADLKYALADPSLKMADPNRFRGKNLPVLDQLTDPPGVKRVYHIQPSLEDPFQPPSIPITVYYAVLERACRSVLLHAPSEAPQIVRGASDEARKHTYNLTIAWYRMGDNCAIPITVMEYTECPYNKSLGVCPIRTQPRWS YYDSFSAVSEDNLGFLMHAPAFETAGTYLRLVKINDWTEITQFILEHRARASCKYALPLRIPPAACLTSKAYQQGVTVDSIGMLPRFIPENQRTVALYSLKIAGWHGPKPPYTSTLLPPELSDTTNATQPELVPEDPEDSALLEDPAGTHHHHHH

## (C) gB2 mRNA-encoded antigen

**gB2 AA sequence (mRNA-encoded antigen):**

<u>MRGGGLICALVVGALVAAVASA</u>APAAPAAPRASGGVAATVAANGGPASRPPPVPSPATTKARKRKTKKPP KRPEATPPPDANATVAAGHATLRAHLREIKVENADAQFYVCPPPTGATVVQFEQPRRCPTRPEGQNYTEGIAVVFKENIAPYKFKATMYYKDVTVSQVWFGHRYSQFMGIFEDRAPVPFEEVIDKINTKGVCRSTAKYVRNNMETTAFHRDDHETDMELKPAKVATRTSRGWHTTDLKYNPSRVEAFHRYGTTVNCIVEEVDARSVYPYDEFVLATGDFVYMSPFYGYREGSHTEHTSYAADRFKQVDGFYARDLTTKARATSPTTRNLLTTPKFTVAWDWVPKRPAVCTMTKWQEVDEMLRAEYGGSFRFSSDAISTTFTTNLTEYSLSRVDLGDCIGRDAREAIDRMFARKYNATHIKVGQPQYYLATGGFLIAYQPLLSNTLAELYVREYMREQDRKPRNATPAPLREAPSANASVERIKTTSSIEFARLQFTYNHIQRHVNDMLGRIAVAWCELQNHELTLWNEARKLNPNAIASATVGRRVSARMLGDVMAVSTCVPVAPDNVIVQNSMRVSSRPGTCYSRPLVSFRYEDQGPLIEGQLGENNELRLTRDALEPCTVGHRRYFIFGGGYVYFEEYAYSHQLSRADVTTVSTFIDLNITMLEDHEFVPLEVYTRHEIKDSGLLDYTEVQRRNQLHDLRFADIDTVIRADANAA

### (D) Recombinant gB2 protein

HSV-2 gB2 protein was purchased from ACROBiosystems (catalog GLB-H52H3). This protein corresponds to the ectodomain of HSV-2 strain 333 gB (UL27) expressed in HEK293 cells and engineered with a C-terminal His_6_ tag.

### Transmission electron microscopy

Samples were applied to carbon-coated 300-mesh Copper EM grid (Electron Microscopy Sciences, Hatfield, PA) that had been glow discharged (25mA, 30 s, Bal-Tec SCD 005 Sputter Coater). Grids were negatively stained using UranyLess EM Stain (Electron Microscopy Sciences) and imaged on a Tecnai G2 Spirit BioTWIN Transmission Electron Microscope (ThermoFisher Scientific, Hillsboro, OR) operated at 80 kV. Images were acquired with a NanoSprint15 MKII camera (AMT Imaging, Woburn, MA) using AMT Capture Engine software (version 7.0.2.5).

### Synthesis of dye-labeled CpG ODN (ODN1826-Cy5)

ODN1826-Cy5 was prepared and characterized by the Oligo Synthesis Facility in the Keck Biotechnology Resource Lab at Yale University. Briefly, ODN1826-Cy5 was synthesized on an ABI 394 DNA/RNA Synthesizer using cyanoethyl phosphoramidite chemistry. The synthesis was initiated using a 3’-Cy5 CPG support (Glen Research, Oak Grove, VA, USA) and UltraMild chemistry utilizing Pac-dA, iPr-Pac-dG, Ac-dC and dT phosphoramdites. The Standard Cap A reagent was replaced with 5% phenoxyacetic anhydride in tetrahydrofuran/pyridine to reduce the possibility of exchange of the iPr-Pac protecting group on the dG moiety with acetate from the standard acetic anhydride capping mix; 4,5-dicyanoimidazole (0.25 M) in acetonitrile was used as the activator. Following deprotection and subsequent drying, ODN1826-Cy5 was purified by high-performance liquid chromatography using reverse-phase techniques. The pooled fractions were dried. Ethanol precipitation was performed to exchange toxic triethylamine (TEA^+^) with sodium (Na^+^) ions.

### In vitro assays

*Chemokine migration Transwell^®^ assay.* Chemotactic activities of free CXCL9 (PeproTech), free CpG ODN (InvivoGen, San Diego, CA, USA), or BEACON (normalized to the indicated CXCL9 equivalents) were quantified using a 96-well Transwell^®^ system. Briefly, splenic CD8^+^ T cells from naïve C57BL/6 WT mice were purified by negative selection (EasySep™; STEMCELL Technologies, Vancouver, BC, Canada), activated for three to five days with plate-bound anti-CD3ε (2 µg/mL) and soluble anti-CD28 (1 µg/mL) in Roswell Park Memorial Institute (RPMI) medium supplemented with 10% fetal bovine serum, 2 mM L-glutamine, 1% penicillin/streptomycin (pen/strep), 25 mM HEPES, and 100 U/mL recombinant (r)IL-2, and maintained in exponential growth. Cells were labeled with Calcein AM (5 µM, 30 min, 37°C), washed twice in migration buffer (RPMI with 0.1% bovine serum albumin), and resuspended at 1.5 x 10^6^ cells/mL. Fibronectin-coated high-throughput screening Transwell^®^-96 3.0 µm inserts (Corning, Product #3387) were equilibrated as per the manufacturer’s instructions. The lower chamber contained 170 µL migration buffer with either free CXCL9 or BEACON. The upper chamber was seeded with 5 × 10^4^ cells in 30 µL. Plates were incubated for 60 min at 37°C and 5% CO₂. Calcein was released from migrated cells via lysis with 10 µL 10% Triton X-100, and fluorescence was assessed using an excitation wavelength of 485 nm and an emission wavelength of 535 nm (Tecan, Mannedorf, Switzerland). Cell numbers were calculated from a per-plate standard curve generated by serially diluting input cells on the receiving plate. All experimental conditions were run in triplicate and repeated in two independent experiments. The parameters followed validated CXCL9/CXCL10 bioassay protocols (ThermoFisher Scientific, Waltham, MA, USA) that were adapted for murine CD8⁺ T cells to ensure consistency across independent experiments.

*Mouse TLR9 reporter assay.* HEK-Blue™ mTLR9 cells (InvivoGen) were seeded at 3 × 10^4^ cells/well in flat-bottom 96-well plates (Corning) in complete HEK-Blue medium and allowed to adhere for 24 h. Cells were then stimulated for 48 h with vehicle (water), CXCL9 (PeproTech), free CpG ODN 1826 (InvivoGen), or BEACON (normalized to the indicated CpG equivalents). Endotoxin units (EU) were measured in the BEACON doses and confirmed to be <0.1 EU/dose (Endosafe Nexgen-PTS, Charles River Laboratories). After incubation, supernatants (20 μL/well) were transferred to a new plate containing Quanti-Blue detection reagent (InvivoGen; 180 μL/well). This reagent provides a colorimetric-based quantification of secreted alkaline phosphatase (SEAP) produced by HEK-Blue™ mTLR9 cells in response to activation of the TLR9–NF-κB/AP-1 signaling pathway. Plates were developed for 60 min at 37°C and read on a microplate reader according to the manufacturer’s instructions. Reporter activity was expressed as fold-change over vehicle control wells included on the same plates.

*BMDC assay.* BMDCs were generated from the femurs and tibias of 8-week-old female C57BL/6 mice as previously described (*60*). Briefly, femoral and tibial BM was cultured in complete medium (RPMI 1640 supplemented with 10% FBS, 1% pen/strep, 1 mM sodium pyruvate, and 1 μM β-mercaptoethanol) with 20 ng/mL granulocyte-macrophage colony-stimulating factor (GM-CSF) for one week. Non-adherent and loosely adherent fractions were collected for downstream screening. Loosely adherent cells were detached via incubation with 5 mM ethylenediaminetetraacetic acid (EDTA) in PBS for 20 minutes at 4°C. All cultures were maintained at 37°C in 5% CO _2_ in a humidified incubator unless otherwise noted. BMDCs were seeded in a 96-well plate (Corning) at 10^5^ cells per well. After 24 hours of culture, the medium was replaced with fresh medium without additional supplements or fresh medium with 100 μL of CXCL9, CpG ODN, BEACON, or water. After 48 hours of incubation, cells were evaluated by flow cytometry with reagents that include a live/dead dye and Fc-blockade (anti-CD16/32) together with lineage and activation markers mentioned in the Antibody list below. The extent of DC maturation was determined by analysis of the expression of the costimulatory receptor CD86 (clone GL1).

### Mouse Necropsy and Histopathology

Female 8-12-week-old C57BL/6 mice were submitted to the Comparative Pathology Research Core (Department of Comparative Medicine, Yale University School of Medicine) for necropsy, comprehensive gross phenotypic analysis with detailed histopathologic analysis of the vagina, as well as analysis of colon, urinary bladder, and lumbar spinal cord tissues of virus infected mice. Pathologists were blind to the experimental group. Mice were euthanized by CO_2_ asphyxiation, weighed, and then exsanguinated by terminal cardiac puncture. Tissues were immersion-fixed in 10% neutral buffered formalin and subsequently trimmed, placed within a cassette, and processed by routine methods. Tissues were sectioned at 5 µm, stained with hematoxylin and eosin (HE), and cover-slipped by routine methods.

Lumbar spine samples were decalcified post fixation using Decal Solution. Vaginal tissues were examined microscopically and scored semi-quantitatively for three parameters: the presence and extent of inflammation, the presence of severity of mucous, and epithelial hyperplasia modified for evaluation of vaginal tissues from previous studies (*61–63*). Briefly, scores included: within normal limits for the tissue, score of 0; trace/scant (1% - 2.5%, score 0.25); less than minimal (2.5 - 5%, score 0.5); minimal (5 - 10%, score 1); mild (10% - 25%, score 2); moderate (25% - 50%, score 3); marked (50% - 75%, score 4); or severe (75% - 100%, score 5). Following the analysis, the groups were decoded. Scores were summed for each mouse to generate a severity of pathology score, after which each group was averaged and the standard deviation calculated. Urinary bladder, colon, and lumbar spinal cord samples were evaluated for the presence of inflammation. Slides were examined using an Olympus BX53 microscope, photographed using an Olympus DP28 camera, and evaluated with Olympus cellSens Standard software 4.1. The images were optimized, and figures prepared using Adobe Photoshop 23.0.

### Quantification of HSV-2 genomes by qPCR

Genomic DNA was extracted using the MagMAX MVP Standard protocol on the KingFisher Flex automated extraction device (ThermoFisher) as per the manufacturer’s instructions. Briefly, samples were incubated with magnetic beads, binding buffer, and proteinase K, then washed and eluted in 75 μL nuclease-free water. Amplification reactions were run on a CFX96 Real-Time PCR System under the following cycling conditions: 95°C for 2 min followed by 40 cycles of 95°C for 15 sec and 60°C for 30 sec (Bio-Rad, Hercules, CA, US). Genome copies were calculated using a five-point, 10-fold standard curve generated with quantified HSV-2 genomic DNA (American Type Culture Collection VR-734DQ; 6.3 × 10^5^ copies/μL).

For detection of HSV-2 genomes in sensory ganglia at later time points, lumbar and thoracic DRG were harvested from mice euthanized at six months post challenge, homogenized manually in 200 µL PBS using a handheld pestle, and stored at -80°C until processing. Genomic DNA was extracted from 200 µL homogenate using the QIAamp® DNA Mini Kit (Qiagen, Germantown, MD, US) with the following modifications. Samples were incubated with 20 µL QIAGEN Protease, lysed with 200 µL Qiagen Buffer AVL (with carrier RNA added according to manufacturer’s instructions) at 56°C for 10 min, then mixed with 230 µL ethanol prior to loading onto columns. HSV-2 genomes were quantified by qPCR targeting HSV-2 gB as described above. For low-input DRG samples, reactions were run in 20 µL total volume with 2.5 µL template, and select samples were re-run with increased template input (5 µL) if amplification was inconsistent across technical replicates. In parallel, amplification of GAPDH was performed as an extraction and input control using a probe-based assay and the following primers (IDT): forward, 5’-CAAATTCAACGGCACAGTCAAG-3’; and reverse, 5’- ACCAGTAGACTCCACGACATAC-3’. The probe, 5’-(FAM)-ATCTTCCAGGAGCGAGACCCCA-(IBlkFQ)-3’ was designed in-house using Primer3 (*64*).

## Antibody list

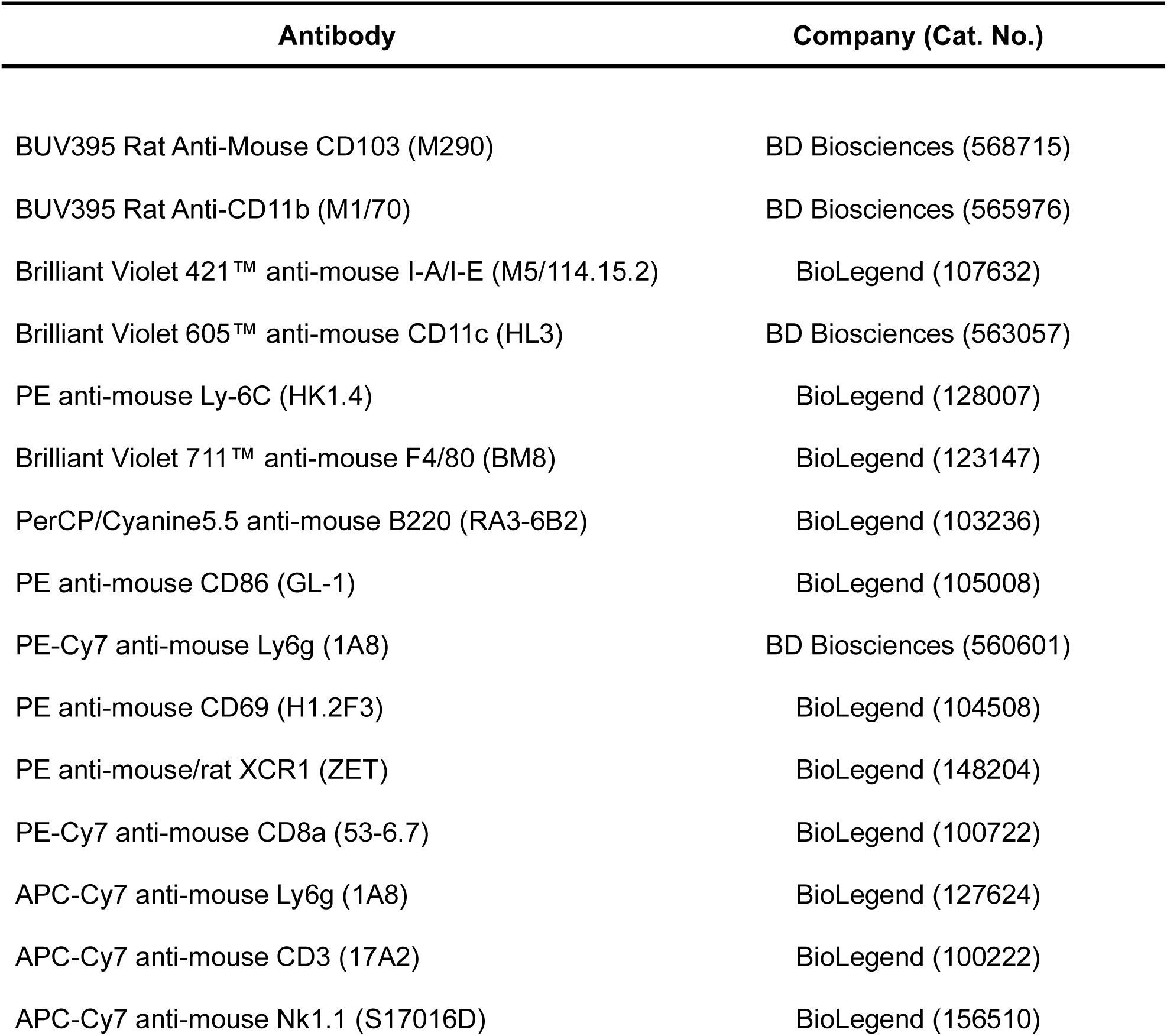
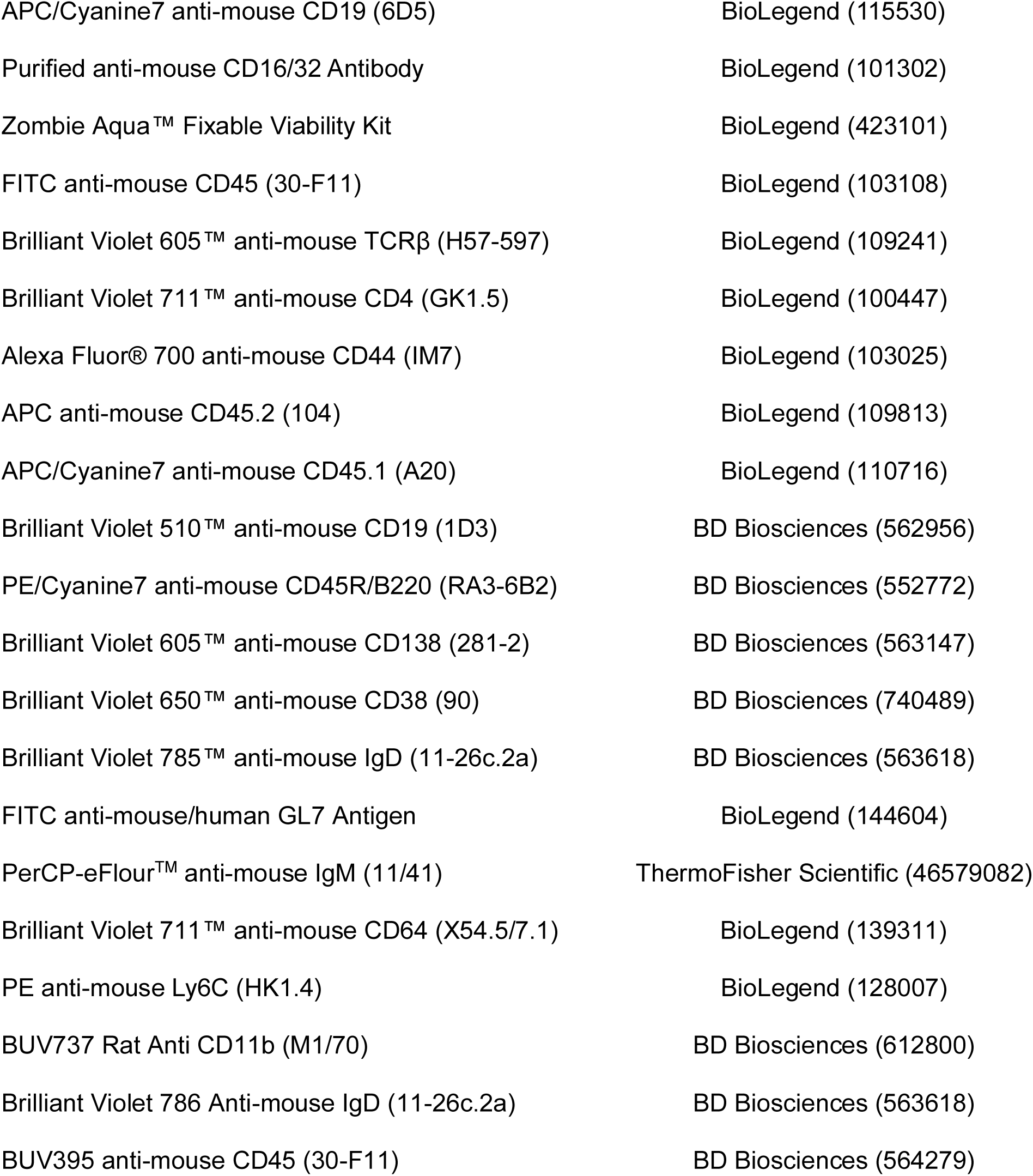

---

**figure S1.**
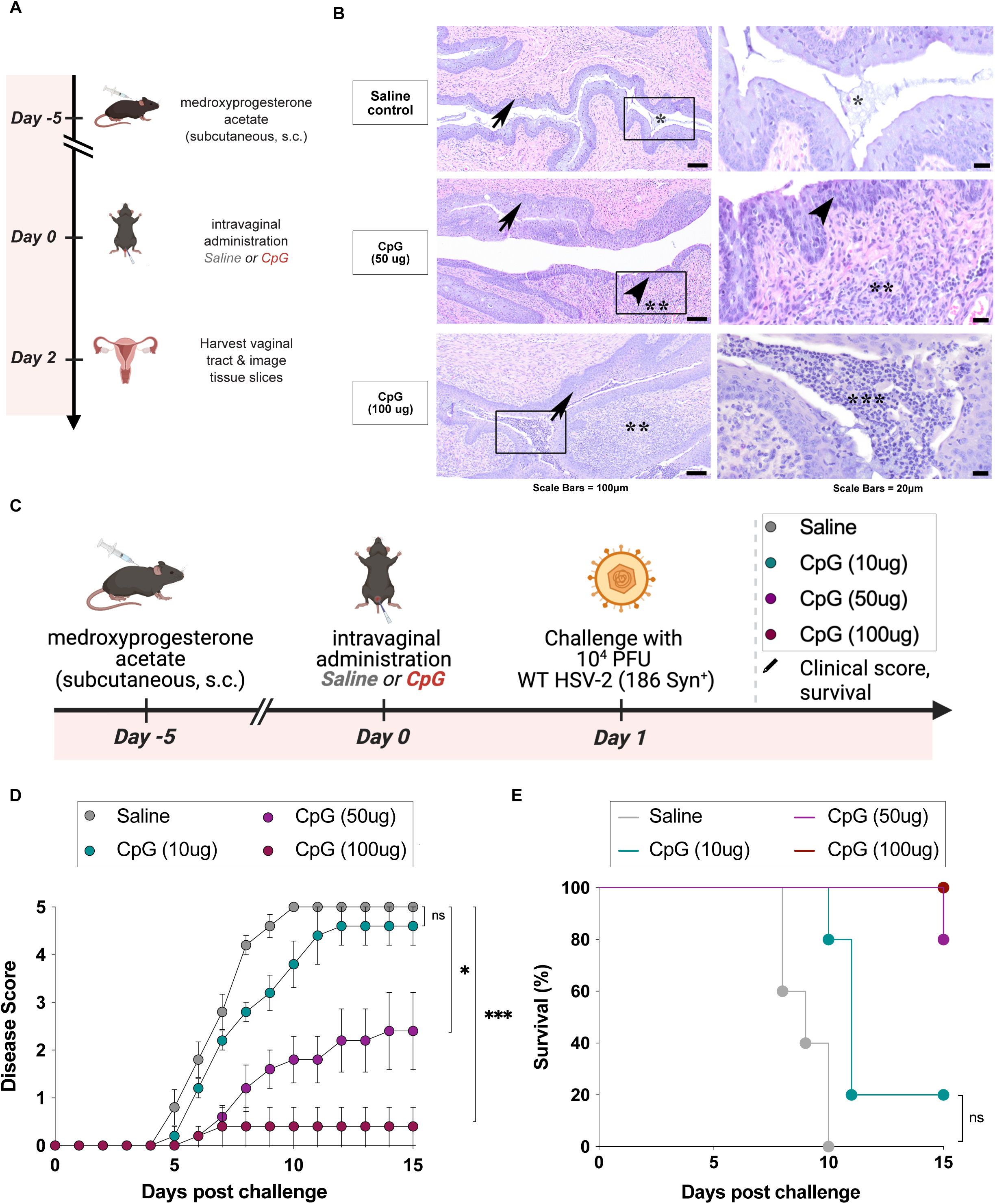
Dose dependent inflammation and protection following intravaginal CpG ODN administration. **(A)** Experimental schematic for CpG ODN tolerability study with vaginal harvest at day 2 post administration. **(B)** Representative HE-stained sections of vaginal tissue from mice treated with saline, 50 µg CpG or 100 µg CpG. **(C)** Schematic for CpG ODN protection study followed by WT HSV-2 challenge. **(D)** Disease scores and **(E)** Kaplan-Meier survival curves in mice treated with saline or CpG ODN (10 ug, 50 ug, or 100 ug) and subsequently challenged with 10^4^ PFU of WT HSV-2 (strain 186 syn^+^). Disease scores were analyzed by two-way repeated-measures ANOVA with Tukey’s multiple comparisons test. Survival differences were assessed by the log-rank (Mantel-Cox) test. *P < 0.05, **P < 0.01, ***P < 0.001, ****P < 0.0001; ns, not significant.

**figure S2.**
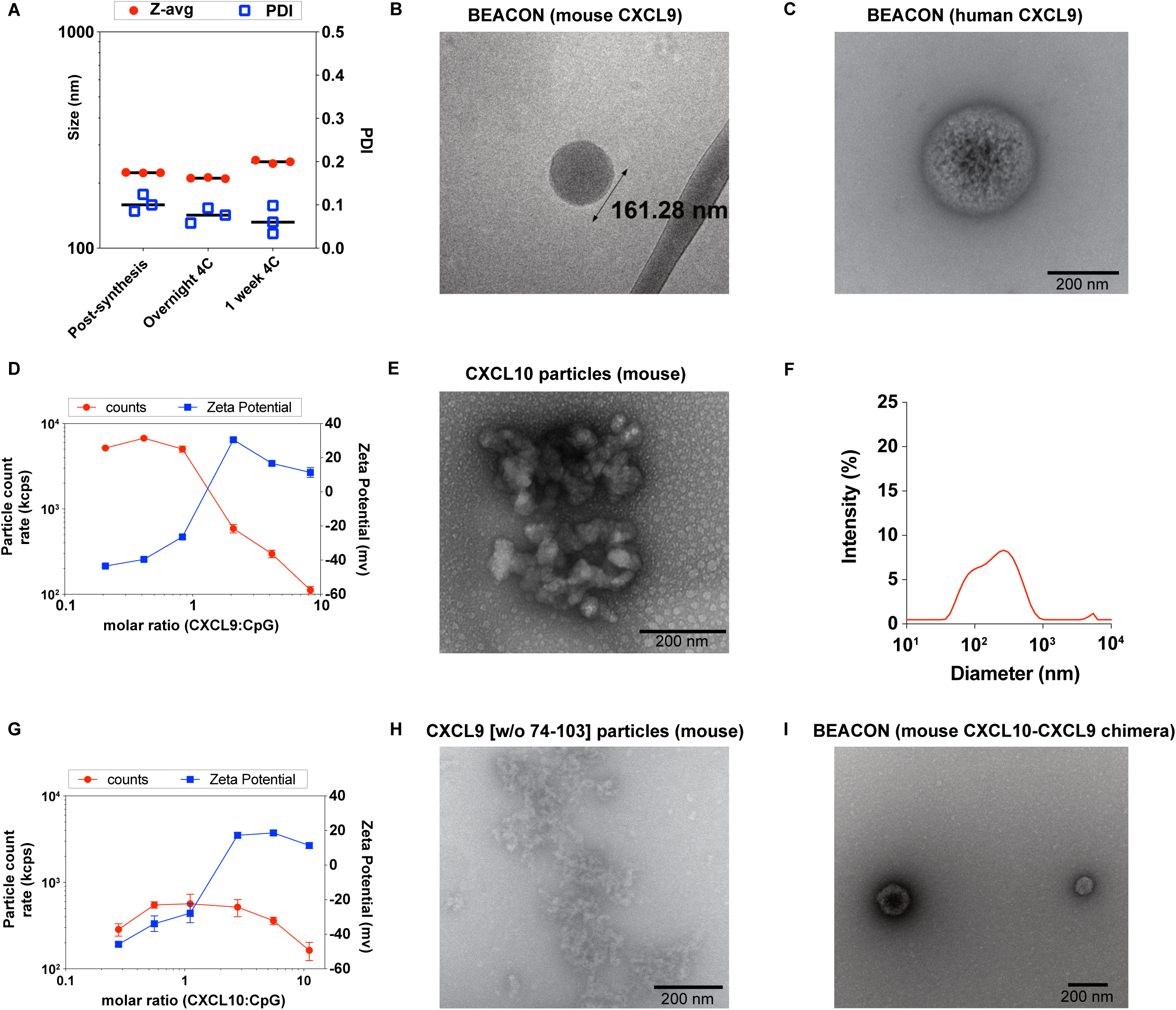
Biophysical characterization of BEACON formulations and comparison to CXCL10-based particles. **(A)** DLS measurements confirming the colloidal stability and uniform size distribution of BEACON (mouse CXCL9) following storage at 4°C for 24 hours or 7 days. Polydispersity index (PDI) values remained low across both time points, indicating retained nanoparticle homogeneity and stability. **(B)** Cryo-transmission electron microscopy (Cryo-TEM) of BEACON (mouse CXCL9) revealing uniform spherical structure. **(C)** Representative negative-stain TEM image of BEACON (human CXCL9) formulated with CpG 1018. **(D)** Particle count and zeta potential measurements for BEACON indicating colloidal stability. **(E-G)** Characterization of CXCL10 particles (mouse): **(E)** TEM image, **(F)** DLS measurements, and **(G)** particle count and zeta potential, revealing poor stability and heterogenous assemblies with CXCL10 (scale bars as indicated). **(H)** TEM image of mouse CXCL9 lacking amino acids 74-103, showing abrogation of nanoparticle formation. **(I)** Negative-stain TEM image demonstrating nanoparticle formation when the mouse CXCL9 C-terminal domain (amino acids 74-103) is appended to mouse CXCL10.

**figure S3.**
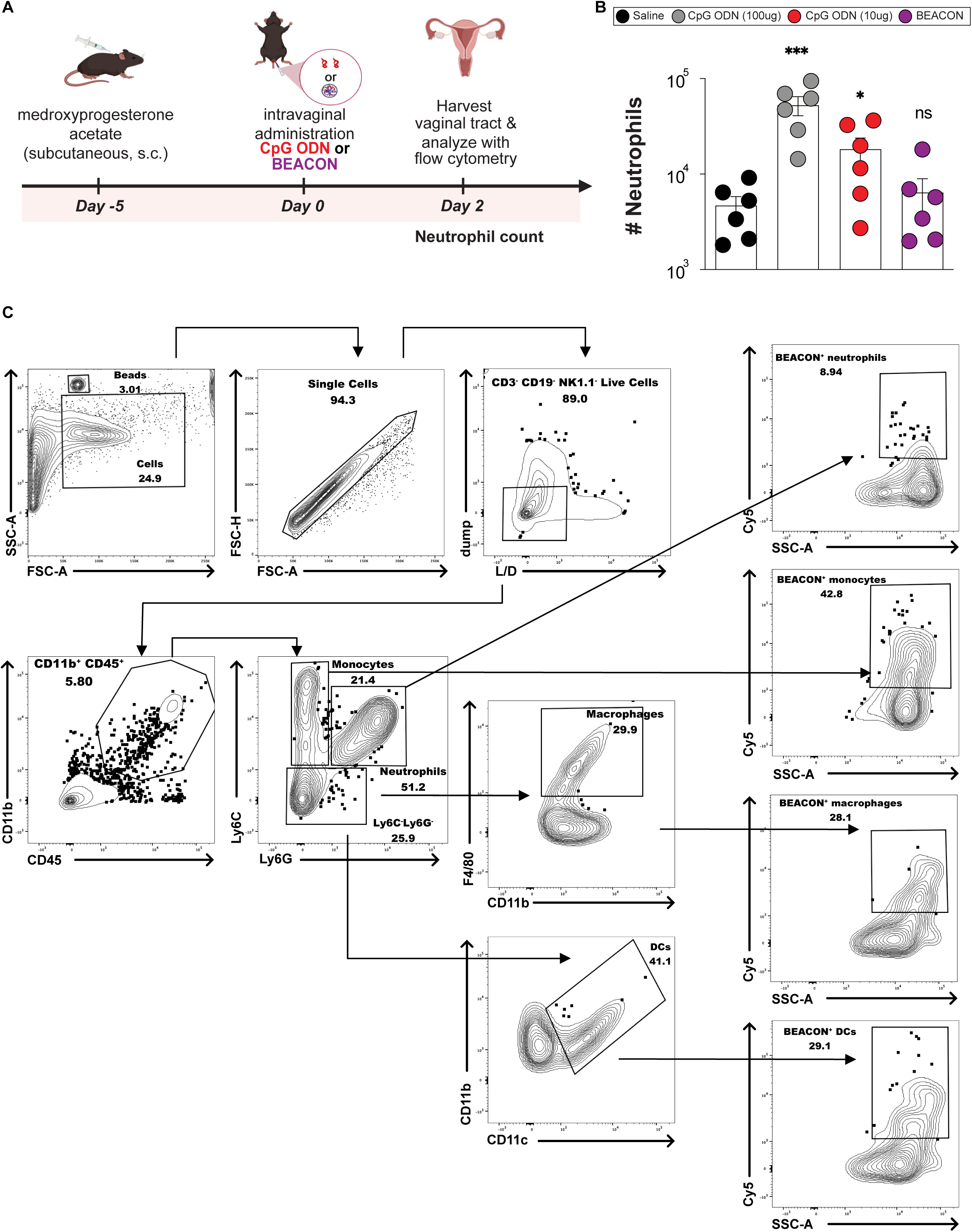
Flow cytometric quantification of vaginal neutrophils and representative gating strategy for identification of APC subsets and Cy5 uptake. **(A)** Schematic for neutrophil quantification following intravaginal administration of saline, CpG ODN, or BEACON, followed by vaginal tissue harvest at day 2 post administration. **(B)** Total vaginal neutrophil counts quantified by flow cytometry (each dot represents one mouse; n indicated by the number of points shown). **(C)** Representative gating strategy for vaginal innate immune cell subsets and quantification of BEACON uptake by neutrophils, monocytes, macrophages, and DCs; dead cells and lymphocyte lineages are excluded. Bars indicate mean ± SEM. Statistical significance was determined by Kruskal-Wallis with Conover-Iman multiple comparisons procedure using Benjamini-Hochberg for controlling false discovery rate (B). *P < 0.05, **P < 0.01, ***P < 0.001, ****P < 0.0001; ns, not significant.

**figure S4.**
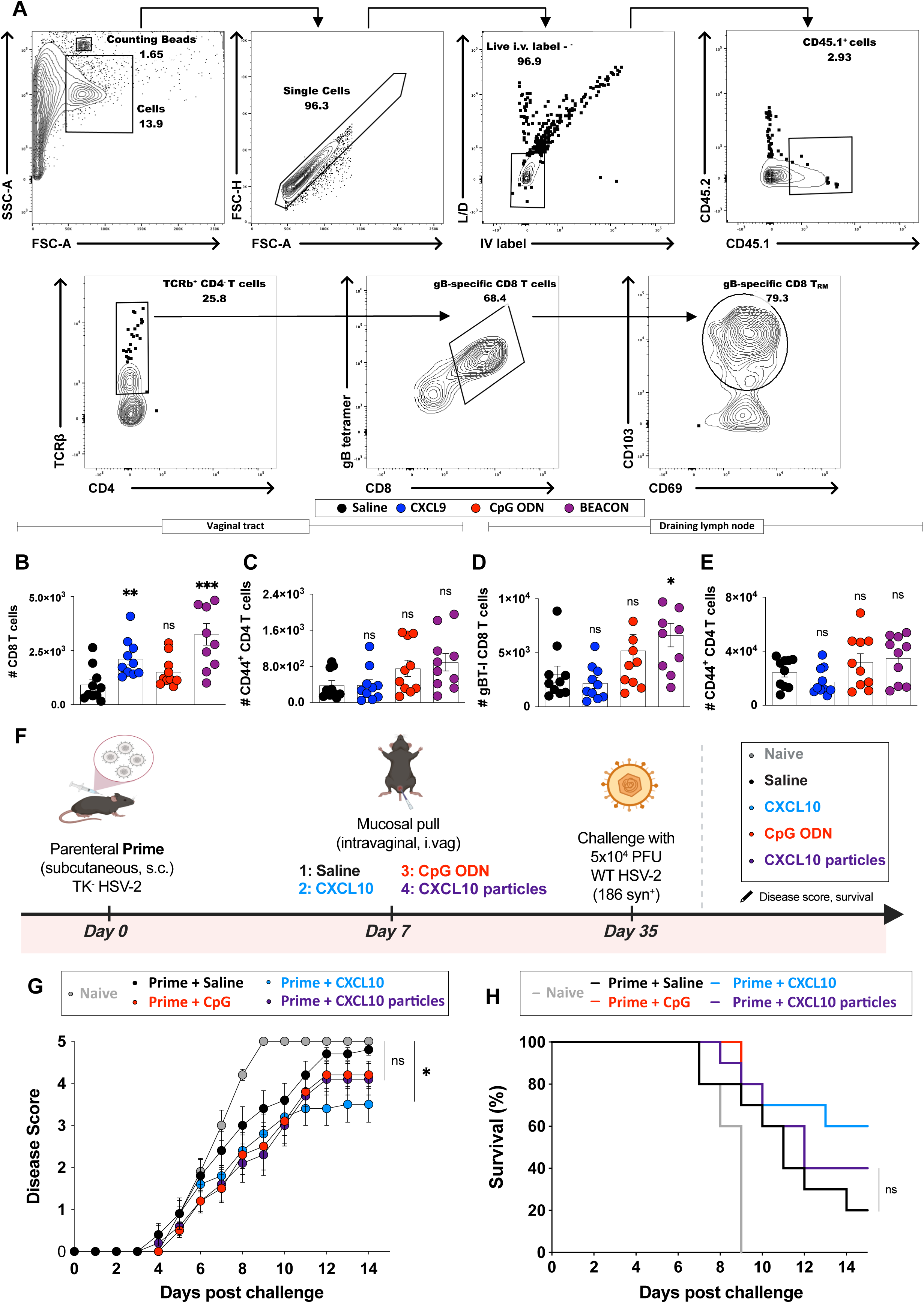
Flow cytometry gating strategy for quantification of vaginal CD8^+^ T_RM_ cells and challenge studies with CXCL10-based particle formulations. **(A)** Representative gating strategy used for quantification of CD8^+^ T_RM_ cells in vaginal tissue. Shown are gates used to identify counting beads, singlets, live cells, exclusion of intravascularly labeled cells, identification of congenic donor cells, gB tetramer^+^ CD8^+^ T cells, and CD69^+^ CD103^+^ T_RM_ cells. **(B-C)** Quantification of total CD8^+^ and CD4^+^ T cells in the vaginal mucosa following the experimental schematic detailed in Fig. 3A. **(D-E)** gBT-I CD8^+^ and CD4^+^ T cell numbers in draining lymph nodes (LNs). **(F)** Schematic of experimental setup for wild-type HSV-2 challenge (strain 186 syn^+^) following mucosal pull with saline, CpG ODN, CXCL10, or CXCL10-containing particles. **(G)** Clinical disease scores and **(H)** Kaplan-Meier survival curves. The results from this experiment indicate that administration of CXCL10-containing particles resulted in no improvement in protection following WT-HSV-2 challenge. Bars in the dot plots indicate mean ± SEM, with each dot representing one mouse. Statistical significance was determined by Kruskal-Wallis with Conover-Iman multiple comparisons procedure using Benjamini-Hochberg for controlling false discovery rate (panels B-E) and by two-way repeated-measures ANOVA with Dunnett’s multiple comparisons test (panel G). Survival differences were assessed by the log-rank (Mantel-Cox) test (panel H). *P < 0.05, **P < 0.01, ***P < 0.001, ****P < 0.0001; ns, not significant.

**figure S5.**
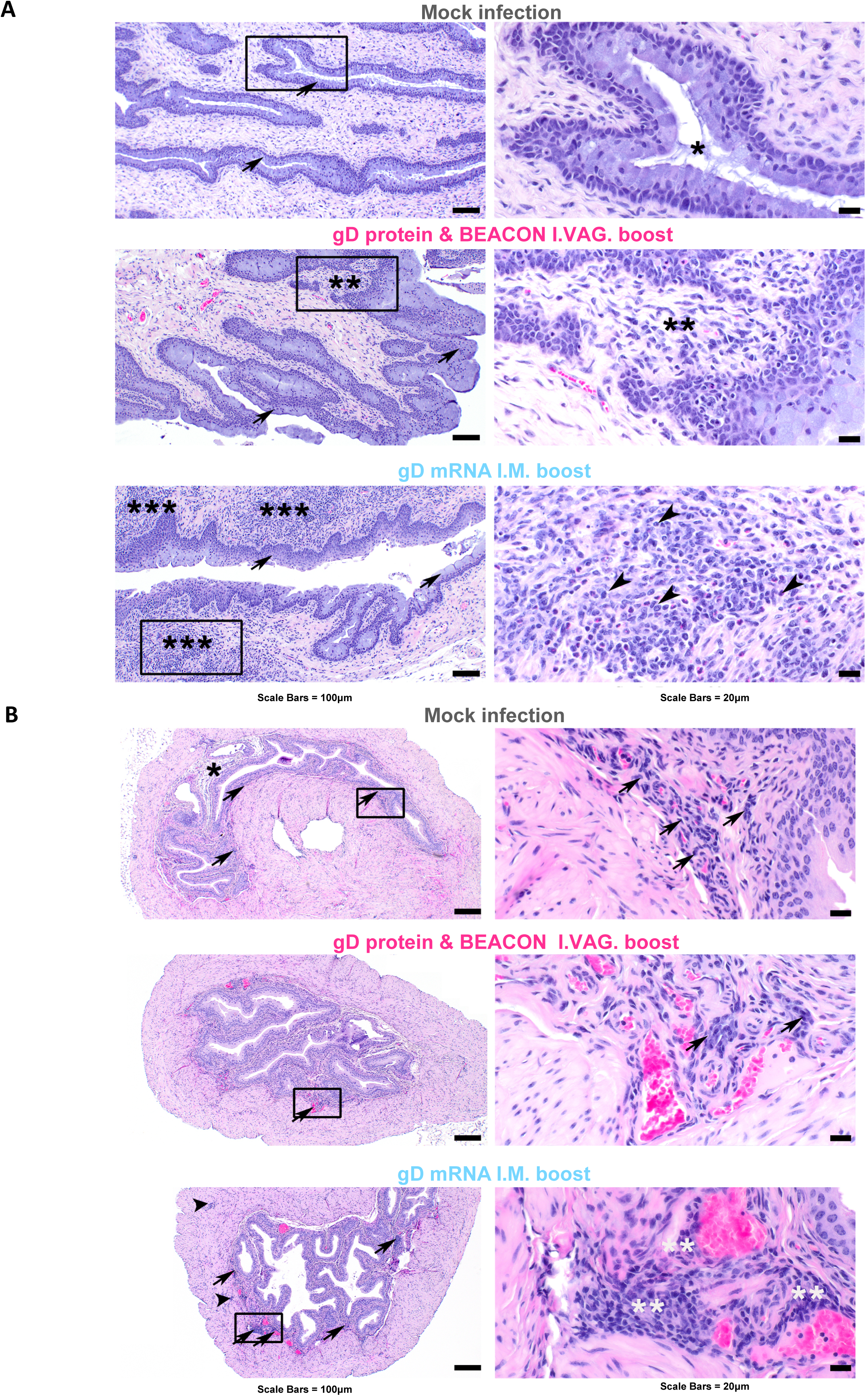
Representative histopathology of vaginal tract and urinary bladder at 6 months post WT HSV-2 virus challenge. **(A)** Representative HE-stained sections vagina from C57BL/6 mice boosted intravaginally with gD protein and BEACON, or intramuscularly with gD mRNA-LNPs taken 6 months post infection with 5 × 10^4^ PFU wild-type HSV-2 (strain 186 syn^+^). While there was no significant inflammatory infiltrate in control or mice boosted intravaginally; there is a marked multifocal submucosal lymphoplasmacytic and neutrophilic/eosinophilic inflammatory infiltrate (**) in the intramuscularly boosted mice. Arrows = mucosal epithelium, Black Arrowheads = neutrophils/eosinophils * = mucous, ** trace/within normal limits inflammation, *** Marked inflammation. **(B)** Representative HE-stained sections of urinary bladder from C57BL/6 mice boosted intravaginally with gD protein and BEACON, or intramuscularly with gD mRNA-LNPs taken 6 months post infection with 5 × 10^4^ PFU wild-type HSV-2 (strain 186 syn^+^). Both control and mice boosted intravaginally have scattered lymphocytes and plasma cells within the laminal propria (arrows), compared to intramuscularly boosted mice that have increased lymphocytes and plasma cells within both the laminal propria and muscularis. Images are representative of mice (n=5) submitted for necropsy within each group.

**figure S6.**
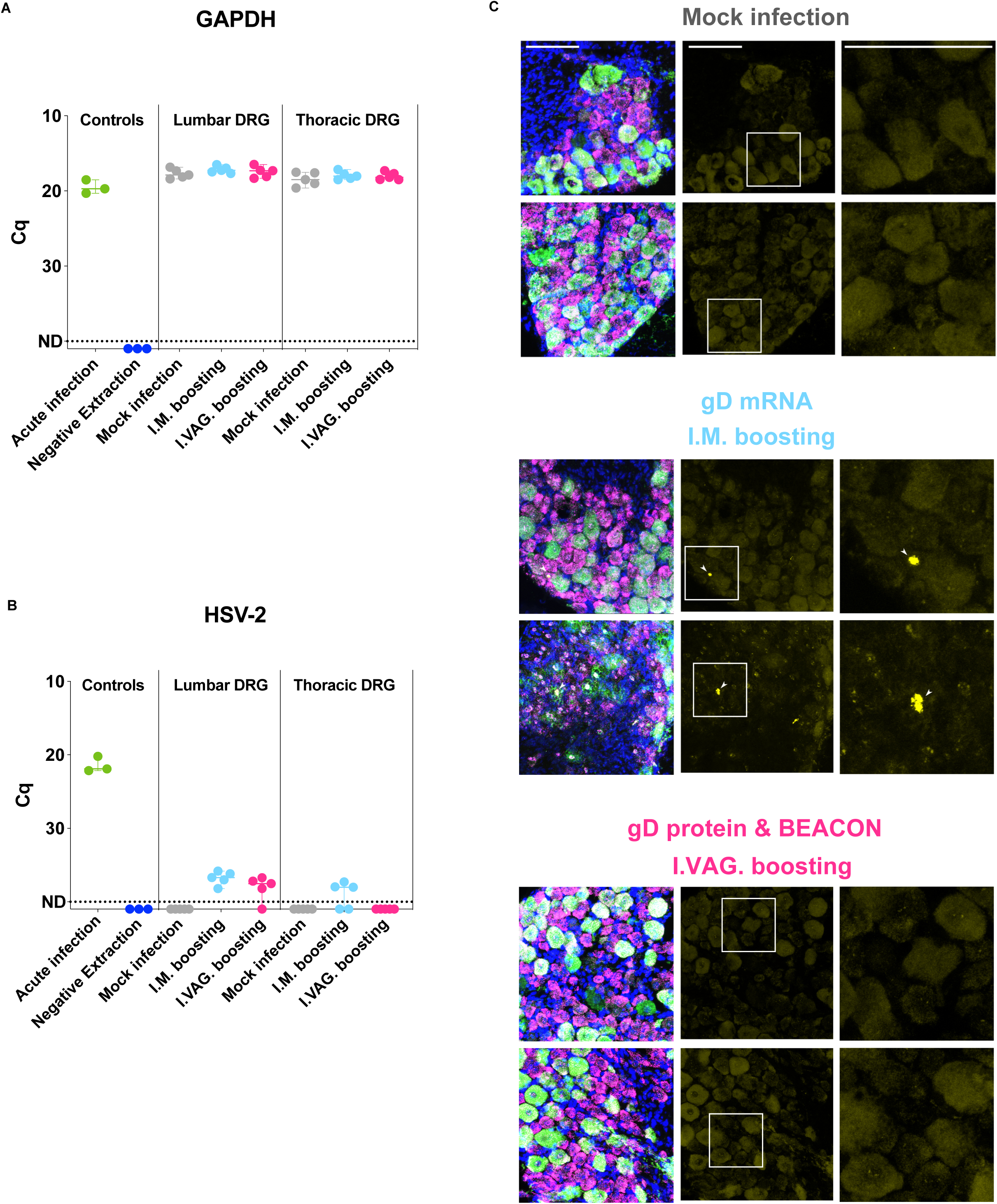
Detection of virus genomes in the DRG at 6 months post challenge by qPCR. **(A)** GAPDH qPCR Cq values from lumbar and thoracic DRG in all groups, including acute infection controls and necropsy endpoints, as indicated in Fig. 5A. **(B)** HSV-2 qPCR Cq values in lumbar and thoracic DRG. **(C)** Additional FISH images of lumbar dorsal root ganglia for each group with probes directed against neurofilament heavy chain (*Nefh*; green), advillin (*Avil*; magenta), or the HSV-2 *LAT* (yellow). Left panels depict the composite overlay of all channels; middle panels show the LAT channel with regions of interest delimited by boxes; right panels show enlarged views of boxed regions. Scale bars = 100 µm and apply to all images.

**figure S7.**
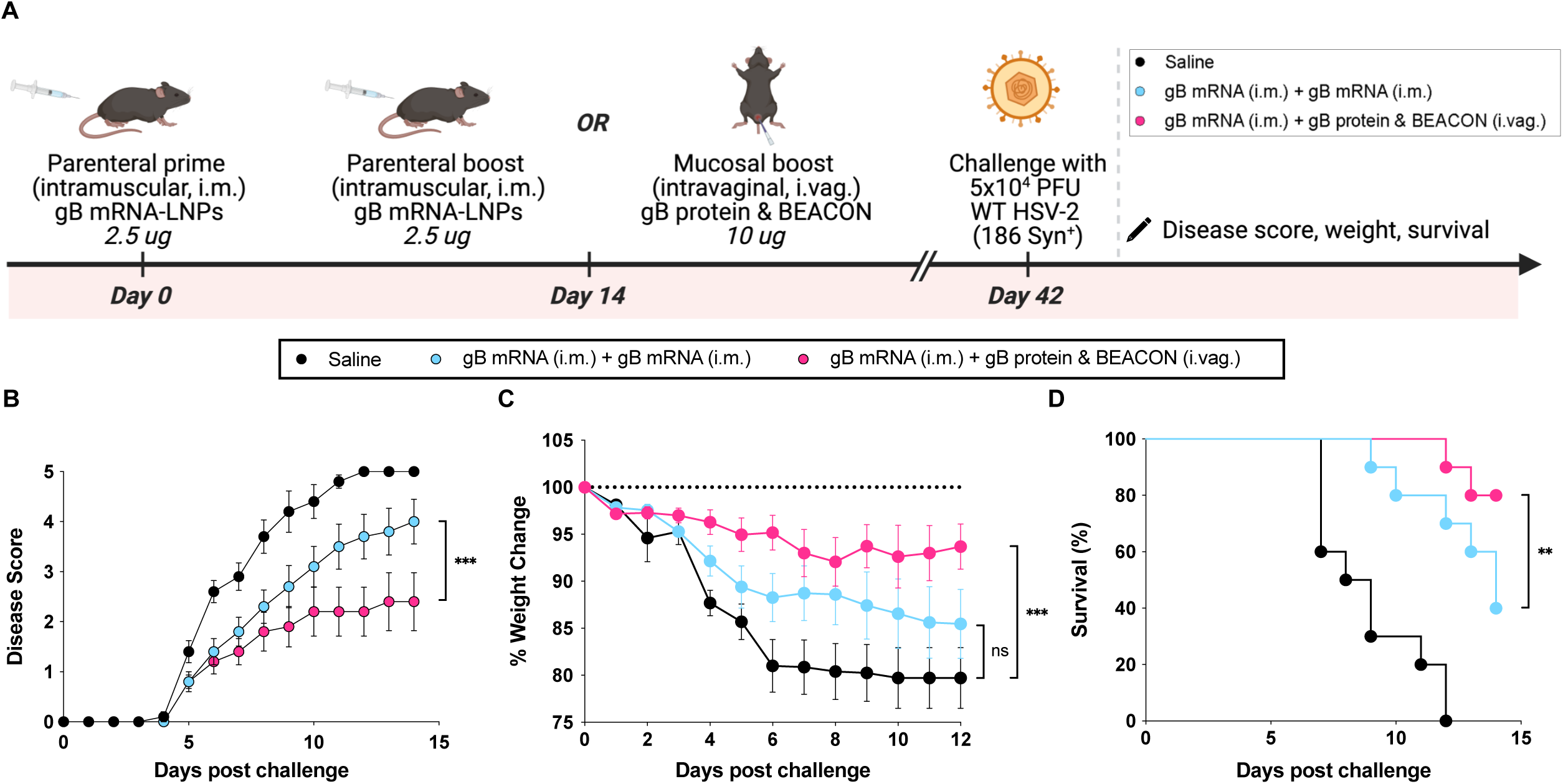
gB-based prime-boost protection study. **(A)** Experimental schematic outlining the gB-based prime-boost immunization strategy. Mice were primed intramuscularly with 2.5 µg gB mRNA-LNPs and four weeks later received either an intramuscular boost with gB mRNA-LNPs or an intravaginal boost with gB protein & BEACON (10 µg). Mice were challenged four weeks post-boost with 5 × 10⁴ PFU HSV-2 (strain 186 syn^+^). **(B)** Clinical disease scores and **(C)** weight loss over time following HSV-2 (strain 186 syn^+^) challenge. **(D)** Kaplan-Meier survival curves indicating superior protection following challenge in the group receiving vaginal gB protein co-administered with BEACON. Disease and weight curves were analyzed by two-way repeated-measures ANOVA with Tukey’s multiple comparisons test. Survival differences were assessed by the log-rank (Mantel-Cox) test. *P < 0.05, **P < 0.01, ***P < 0.001, ****P < 0.0001; ns, not significant.

**figure S8.**
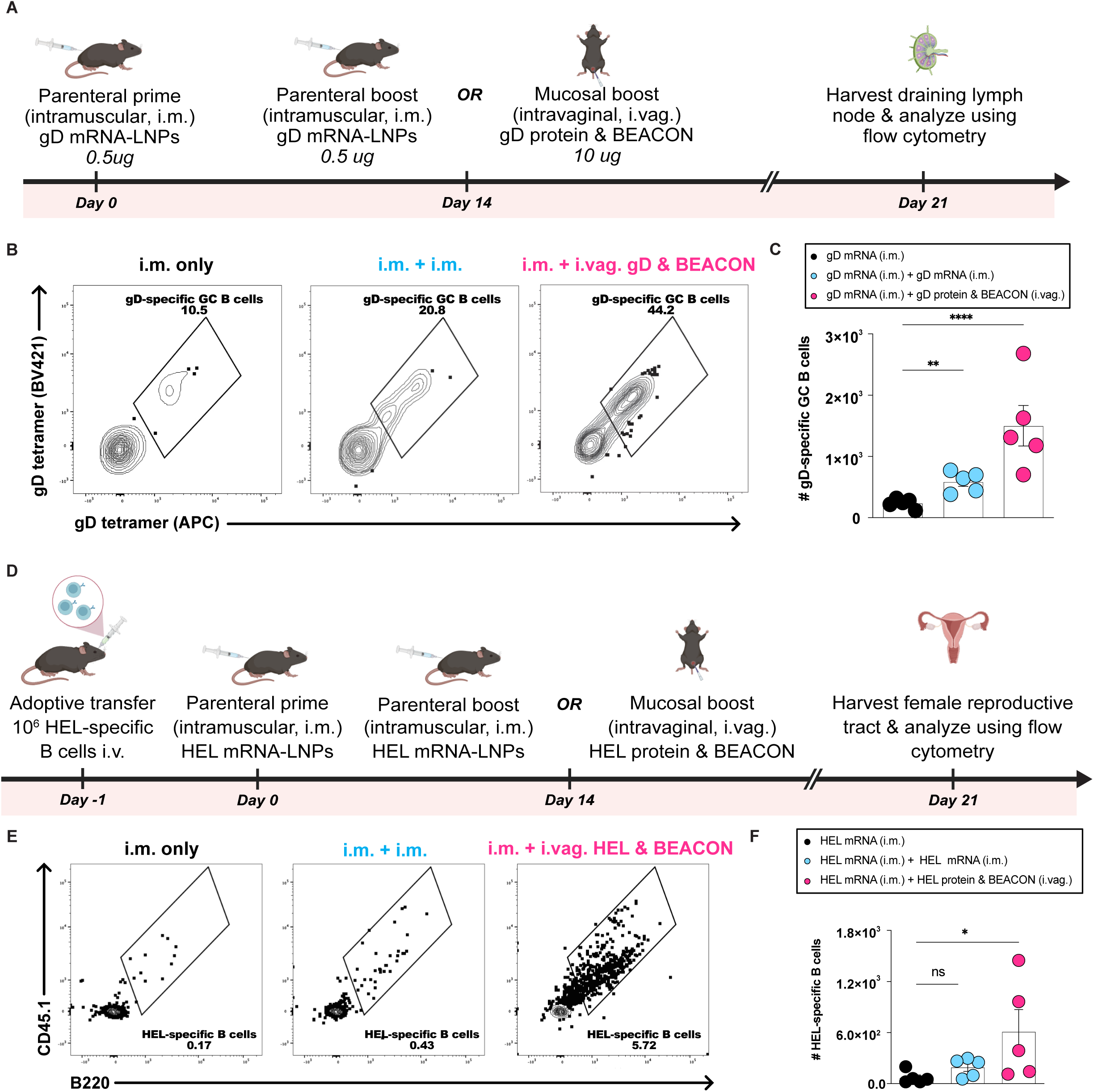
Analysis of B cell responses following mucosal boosting with BEACON. **(A)** Schematic for the analysis of gD germinal center B cells in draining LNs after boosting. **(B)** Representative gating for gD-specific germinal center B cells using gD tetramers. **(C)** Quantification of gD-specific germinal center B cell numbers in draining LNs. **(D)** Schematic for HEL B cell adoptive transfer experiments to assess antigen-specific B cell recruitment to the female reproductive tract (FRT). **(E)** Representative flow cytometry plots for HEL-specific B cells in the FRT. **(F)** Quantification of HEL-specific B cell numbers in the FRT. Each dot represents data from a single mouse. Bars indicate mean ± SEM. Statistical significance was determined by Kruskal-Wallis with Conover-Iman multiple comparisons procedure using Benjamini-Hochberg for controlling false discovery rate (panels C, F). *P < 0.05, **P < 0.01, ***P < 0.001, ****P < 0.0001; ns, not significant.

**figure S9.**
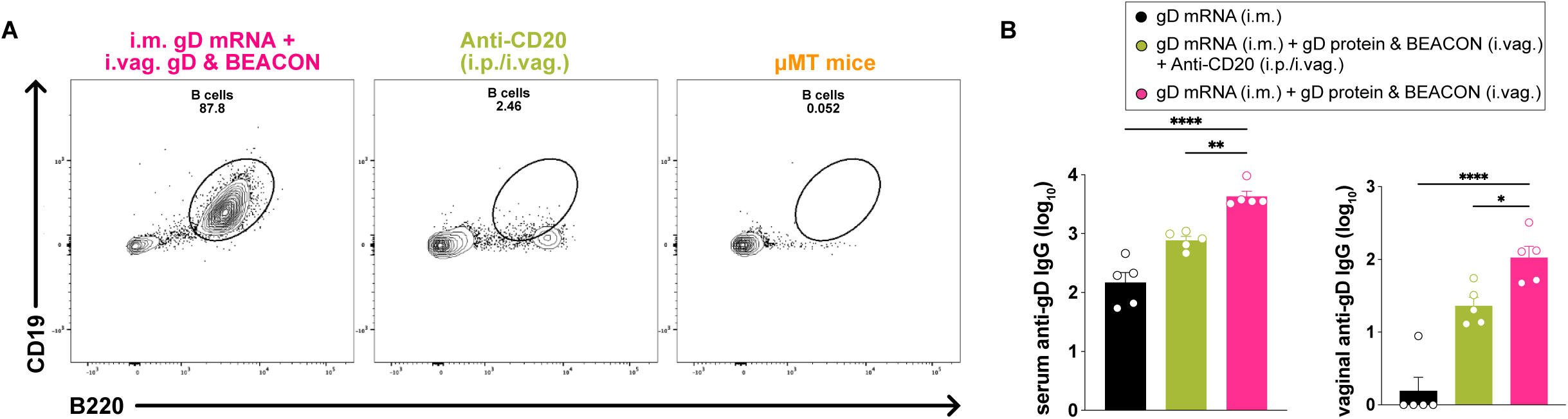
Validation of B cell depletion to delineate the role of protective humoral responses following BEACON-based vaginal boosting. **(A)** Flow cytometric analysis confirming partial depletion of CD19^+^ B220^+^ B cells following anti-CD20 treatment. **(B)** gD-specific serum and vaginal IgG titers in isotype control (pink), anti-CD20-treated (green), and primed-only (black) mice. The results indicate that disease severity correlates inversely with serum and vaginal antibody titers. Bars indicate mean ± SEM. Statistical significance was determined by Kruskal-Wallis with Conover-Iman multiple comparisons procedure using Benjamini-Hochberg for controlling false discovery rate (panel B). *P < 0.05, **P < 0.01, ***P < 0.001, ****P < 0.0001; ns, not significant.

**figure S10.**
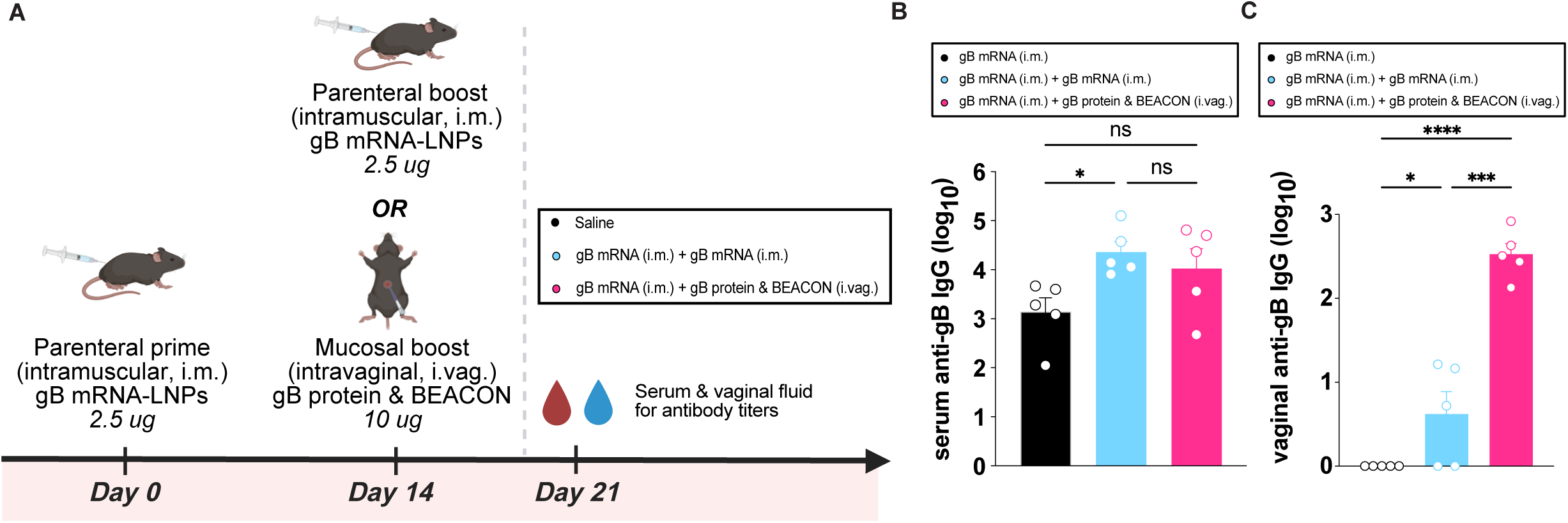
Antibody responses following mucosal boosting with HSV gB protein and BEACON. **(A)** Schematic for evaluation of gB antibody responses to prime and boost regimens. **(B)** Serum anti-gB IgG titers. **(C)** Vaginal anti-gB IgG titers. Bars indicate mean ± SEM. Statistical significance was determined by Kruskal-Wallis with Conover-Iman multiple comparisons procedure using Benjamini-Hochberg for controlling false discovery (panels B and C). *P < 0.05, **P < 0.01, ***P < 0.001, ****P < 0.0001; ns, not significant.

**figure S11.**
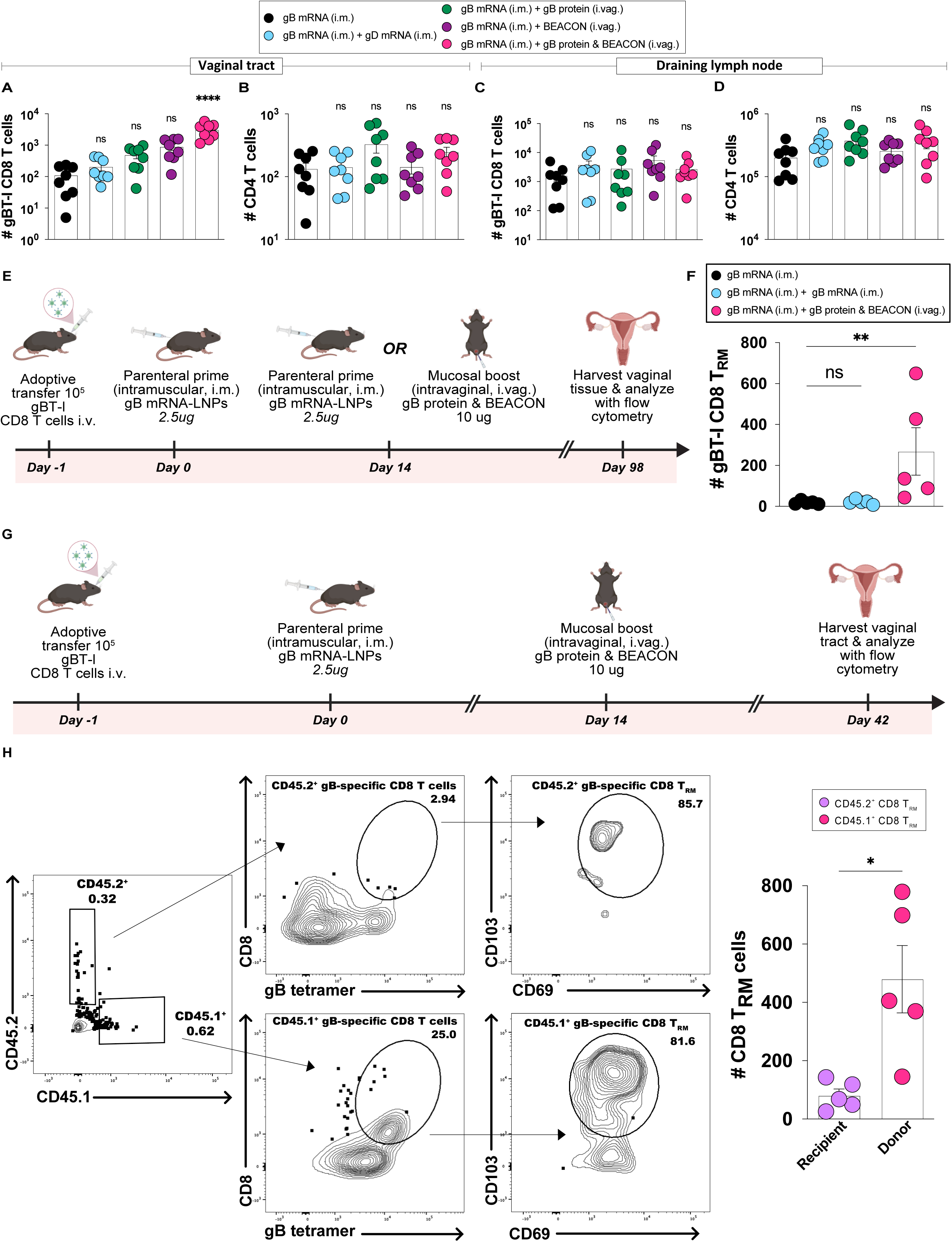
Total vaginal CD8^+^ and CD4^+^ T cell counts and durability of gB-specific T_RM_ responses following mucosal boosting with gB protein and BEACON. C57BL/6 mice received 10⁵ congenically marked gBT-I CD8^+^ T cells intravenously (day -1) followed by intramuscular priming with 2.5 µg gB mRNA-LNPs (day 0). On day 14, mice received one of four boosts: (i) intramuscular gB mRNA-LNPs (2.5 µg); (ii) intravaginal gB protein alone (10 µg); (iii) intravaginal BEACON alone; or (iv) intravaginal co-delivery of gB protein (10 µg) with BEACON. Vaginal tissues and iliac draining LNs were harvested 4 weeks post-boost (day 42) and analyzed by flow cytometry using counting beads. **(A)** gBT-I CD8^+^ T cell counts, and **(B)** CD4^+^ T cell counts in the vaginal tract. **(C)** gBT-I CD8^+^ T cell counts, and **(D)** total CD4^+^ T cell counts in the draining iliac LNs. **(E)** Experimental schematic for durability analysis following intravaginal boosting with gB protein and BEACON versus intramuscular boosting with gB mRNA-LNPs; vaginal tissues were harvested at day 98 post-boost. **(F)** Number of vaginal CD8^+^ T_RM_ cells (CD69^+^CD103^+^gBT-I) at day 98 in response to each of the regimens indicated. (Each dot represents data from a single mouse; n is indicated by the number of points shown). **(G)** Experimental schematic and **(H)** representative flow plots showing discrimination between congenic populations and CD69^+^CD103^+^ T_RM_ gating for CD45.2^+^ and CD45.1^+^ gB-specific CD8^+^ T cell populations, with corresponding quantification. For dot plots, bars indicate mean ± SEM. Each dot represents data from a single mouse. Statistical significance was determined by Kruskal-Wallis with Conover-Iman multiple comparisons procedure using Benjamini-Hochberg for controlling false discovery rate (panels A-F) and by two-tailed Student’s t-test for donor versus recipient comparisons (panel H). *P < 0.05, **P < 0.01, ***P < 0.001, ****P < 0.0001; ns, not significant.

**figure S12.**
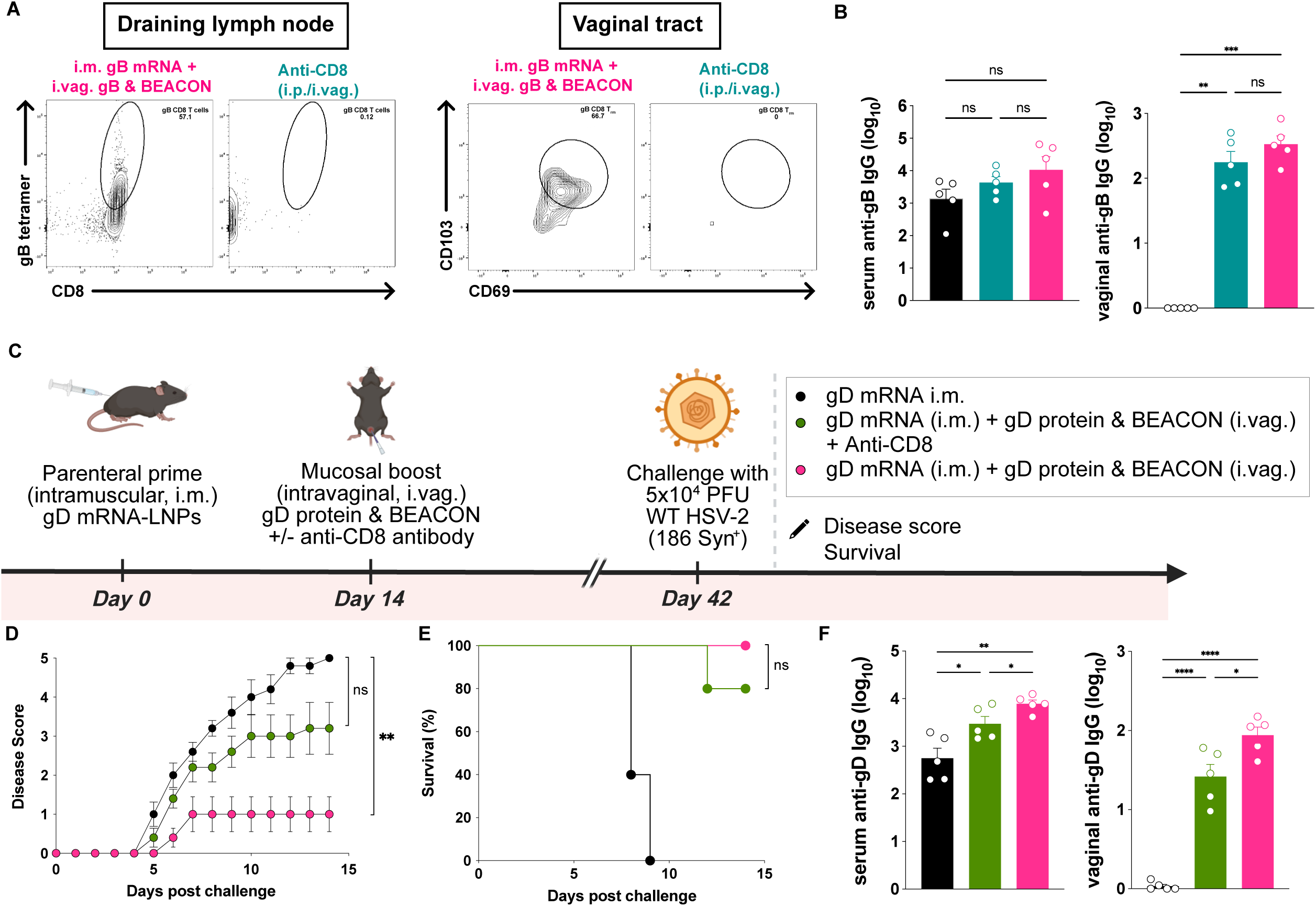
CD8^+^ T cells are required for the induction of protective mucosal immunity following BEACON-based vaginal boosting. **(A)** Representative FACs plots validating CD8^+^ T cell depletion in the LNs and vaginal tract following anti-CD8 antibody administration. **(B)** Quantification of gB-specific IgG in serum and vaginal washes of CD8-depleted mice. **(C)** Experimental schematic of immunization and depletion strategy: mice were primed intramuscularly with gD mRNA-LNPs, boosted intravaginally with gD protein & BEACON, and treated with anti-CD8 antibody prior to HSV-2 challenge. **(D)** Disease scores and **(E)** Kaplan-Meier survival curves showing increased pathology in CD8-depleted mice relative to isotype controls. **(F)** Quantification of gD-specific IgG in serum and vaginal washes of CD8-depleted mice. Bars indicate mean ± SEM. Antibody titers and viral genome comparisons were analyzed by Kruskal-Wallis with Conover-Iman multiple comparisons procedure using Benjamini-Hochberg for controlling false discovery (B, F). Disease scores were analyzed by two-way repeated-measures ANOVA with Tukey’s multiple comparisons test (D). Survival differences were assessed by the log-rank (Mantel–Cox) test (E). *P < 0.05, **P < 0.01, ***P < 0.001, ****P < 0.0001; ns, not significant.

